# Parasitic Vectors in Aquaculture: *Neobenedenia girellae* and Leeches as Potential Transmission Agents for *Trypanosoma carassii spectrum* and Pathogenic bacteria

**DOI:** 10.64898/2026.08.23.746442

**Authors:** Jinsong Chen, Jingyu Zhuang, Xintao Li, Musen Lin, Qianyu Lu, Ning Yan, De-Hua Lai, Shengfeng Huang

**Affiliations:** State Key Laboratory of Biocontrol and School of Life Sciences, Guangdong Provincial Key Laboratory for Aquatic Economic Animals, Guangdong Key Laboratory of Pharmaceutical Functional Genes, Sun Yat-sen University, Guangzhou 510275, China

**Keywords:** Fish Parasites, Vector, leech, trypanosome, *Neobenedenia girellae*, pathogenic bacterium

## Abstract

Parasitic infections pose multifaceted threats to farmed fish, extending beyond direct pathogenicity to facilitate infections of bacteria, viruses, and microparasites. This synergistic interaction often leads to co-infections that significantly exacerbate disease outbreaks and mortality, presenting a severe challenge to aquaculture sustainability. Recently, a novel trypanosomiasis caused by the *Trypanosoma carassii spectrum* has emerged in cage-cultured *Larimichthys crocea* along the southeast coast of China, resulting in widespread prevalence and high mortality rates. Although this pathogen is hypothesized to originate from freshwater fish, its transmission route in marine environments has remained elusive. In this study, we investigated potential vectors and intermediate hosts of *T. carassii spectrum*, including leeches and monogenean in natural marine settings, and simulated transmission pathways using an established laboratory model involving *T. carassii spectrum*, *Micropterus salmoides* and the leech *Poecilobdella manillensis*.

First, our field surveys in the coast of Ningde, Fujian Province, revealed a nearly 100% co-infection rate of *T. carassii spectrum* and the monogenean *Neobenedenia girellae* in diseased juvenile *L. crocea*. PCR analysis detected *T. carassii spectrum* traces in some *N. girellae* specimens, and subsequent experiments confirmed that *N. girellae* ingests the trypanosome while feeding on host blood. Furthermore, bacterial co-pathogens, such as *Vibrio harveyi*, were also detected within *N. girellae*. We also document two fatal leech infestations: *Zeylanicobdella arugamensis* in hybrid groupers (*Epinephelus moara* ♀ X *Epinephelus lanceolatus ♂*) in Zhangpu, and *Limnotrachelobdella okae* in *E. lanceolatus* and *E. fuscoguttatus* in Raoping. These leeches tested negative for trypanosomes but carried pathogenic bacteria that co-infected the host fish; nonetheless, they are established vectors for trypanosome transmission.

In a laboratory cohabitation model simulating *T. carassii spectrum* transmission, infected *M. salmoides* were housed with healthy conspecifics under three conditions: Group A (with the leech *P. manillensis*), Group B (no leeches), and Group C (no leeches, with physical separation between infected and healthy fish). After 14 days, blood smear microscopy and PCR analysis revealed infection rates in healthy fish of 58.33% in Group A, 40.00% in Group B, and 0% in Group C. Conclusively, *T. carassii spectrum* can be transmitted via leeches (with higher efficiency) and may also spread through direct contact under high-density aquaculture conditions, whereas *N. girellae* may act as an incidental vector, further research is warranted to clarify transmission dynamics in natural marine ecosystems. Additionally, our findings highlight the role of ectoparasites, including *N. girellae* and leeches, as potential reservoirs and vectors for bacterial pathogens of fish. In high-density intensive aquaculture, this vectorial capacity transforms parasites from primary pathogens into key drivers of polymicrobial disease outbreaks.

## 1. Introduction

Parasitic diseases pose a persistent threat to aquaculture, causing direct economic losses and indirect yield reductions. Intensive farming conditions facilitate parasite transmission; notably, outbreaks of the monogenean *Gyrodactylus salaris* in Norwegian *Salmo salar* farming have led to mass mortality and stringent stocking restrictions (Peeler et al., 2006). In intensive freshwater aquaculture, infections with *Ichthyophthirius multifiliis* frequently result in high mortality rates and substantially increased therapeutic costs (Abu-Elala et al., 2021). Marine cage aquaculture also faces significant challenges, where sea lice infections impair growth performance and induce mortality in *S. salar* (Bricknell et al., 2006). Infection with *Benedenia humboldti* substantially compromises the health of *Seriola lalandi*, resulting in reduced feed conversion efficiency, growth retardation, and elevated culling rates (Baeza et al., 2019). Furthermore, outbreaks of the ciliate *Cryptocaryon irritans*, a common pathogen in marine fish culture, significantly reduce aquaculture production (Li et al., 2022). Thus, parasitic diseases have become a critical constraint on the sustainable development of aquaculture. The pathogenicity of parasites extends beyond direct tissue damage to include complex immunosuppressive mechanisms. Monogeneans, such as *Neobenedenia* spp. and *Gyrodactylus* spp., cause epithelial erosion of the skin and gills through attachment and feeding, leading to osmoregulatory imbalance and hypoxia (Ogawa, 2015). Skin ulceration resulting from sea lice blood-feeding not only induces stress responses but also provides an entry point for secondary bacterial infections (Velloso and Pereira, 2010), tomonts formed by *I. multifiliis* on the gills, accompanied by epithelial hyperplasia, significantly impair respiratory gas exchange efficiency in fish (Xu et al., 2012). More insidiously, various helminths secrete immunomodulatory molecules that suppress both mucosal and systemic immune responses in the host, thereby increasing susceptibility to secondary infections (Perez-Cordon et al., 2014). Blood-borne trypanosomes cause anemia and immune suppression, significantly increasing host susceptibility to bacterial sepsis (KIRMSE, 1980).

In intensive aquaculture systems, co-infections involving parasites, bacteria, and viruses are common and frequently exert pathogenic effects. We hypothesize that certain macroparasites, particularly hematophagous species such as sea lice, leeches and monogeneans, may act as mechanical vectors or biological vectors for other pathogens, thereby serving as primary drivers of disease outbreaks. Studies indicate that sea lice infestations are frequently accompanied by secondary infections with *Vibrio alginolyticus* and *Aeromonas salmonicida* (Okon et al., 2023). Following mucosal damage induced by *I. multifiliis*, infection rates of opportunistic pathogens such as *Aeromonas hydrophila* increase significantly (Abu-Elala et al., 2021). Monogenean infections are frequently co-detected with *Streptococcus* and *Vibrio* species, leading to significantly increased mortality rates (Ogawa, 2015). Furthermore, hematophagous ectoparasites (e.g., sea lice and leeches) have been implicated in experimental and epidemiological studies as mechanical vectors or facilitators of the inter-population transmission of viruses (Infectious Salmon Anaemia Virus) and bacteria (Barker et al., 2019; Oelckers et al., 2014).

In recent years, trypanosomiasis caused by a novel *Trypanosoma carassii spectrum* has emerged in large yellow croaker (*Larimichthys crocea*) along China’s southeastern coast, causing substantial economic losses to the aquaculture industry (Qin et al., 2025; Wang et al., 2025). Although this pathogen is likely of freshwater fish origin, its transmission route in *L. crocea* remains elucidated. Residing in the host’s blood, fish trypanosomes are most commonly spread through hematophagous intermediaries, particularly leeches, that breach the integumentary barrier (Fermino et al., 2015). The presence of specific symbiotic microbiota in hematophagous leeches, combined with their blood-feeding behavior, provides the biological basis for the potential transmission of hemoprotozoan parasites (Manzano-Marín et al., 2015; Perkins et al., 2005). It is noteworthy that marine leech infestations have also surged in southeastern China recently, albeit in fish species other than *L. crocea* (Che et al., 2026). Whether these leeches serve as vectors for the trypanosomes infecting *L. crocea* remains to be elucidated.

This study conducted field surveys and detection of *T. carassii spectrum* infections in *L. crocea* and potential vectors (leeches and monogenean) along China’s southeastern coast. Furthermore, we employed an established laboratory model involving *T. carassii spectrum*, largemouth bass (*Micropterus salmoides*) and the leech *Poecilobdella manillensis* to simulate transmission pathways. These efforts aim to elucidate transmission mechanisms of *T. carassii spectrum*, enrich the theoretical framework of parasite-mediated pathogen transmission, and provide a scientific basis for integrated disease control strategies in aquaculture.

## 2. Materials and methods

### 2.1 Field survey

Field surveys were conducted to investigate three distinct disease outbreaks in coastal marine aquaculture facilities in Southeast China. In August 2025, a trypanosomiasis outbreak occurred in net-cage cultured *L. crocea* in Ningde City, Fujian Province (26.5904°N, 119.8637°E). Subsequently, epizootics of fish leech infestation were reported in grouper farms in Zhangpu County, Fujian Province (24.1312°N, 117.9425°E) in November 2025, and in Raoping County, Guangdong Province (23.5675°N, 117.0816°E) in January 2026. Our team performed on-site investigations for each case, documenting cultivation models, environmental parameters (water temperature and salinity), clinical signs, fish specifications, mortality rates, and parasite infection intensities. Representative fish exhibiting typical symptoms were selected for the collection and identification of parasites and associated bacterial pathogens.

### 2.2 Collection of Parasites

Collection and Preservation of Trypanosomes: Blood samples were collected from the caudal vein of clinically affected fish in the field. Syringes and centrifuge tubes were pre-rinsed with 1 mg/mL sodium heparin (CAS No. 9041-08-1, Solarbio, China) to prevent coagulation. Glycerol was added to the anticoagulated blood to a final concentration of 10%, and the mixture was thoroughly homogenized and preserved on dry ice. Upon arrival at the laboratory, approximately 50 μL of the sample was used directly for microscopic examination, while the remaining volume was reserved for genomic DNA extraction.

Collection and Fixation of benedeniid monogeneans: Adult benedeniid monogeneans were collected from the skin and eyes of clinically affected fish. To facilitate detachment, infected fish were immersed in freshwater, causing the monogeneans to turn white and detach due to osmotic stress. The detached parasites were collected and either fixed in 70% ethanol for long-term storage or snap-frozen in liquid nitrogen for subsequent genomic DNA extraction. Additionally, 3-5 live specimens were fixed in ammonium picrate solution and flattened for morphological identification and microscopic examination.

Collection and Preservation of Fish Leeches Fish leeches, which are macroscopic ectoparasites visible to the naked eye, were collected from the skin of infected groupers using forceps or pipettes. To facilitate detachment from the host epidermis, a small amount of alcohol was applied directly to the attachment sites of the suckers. A subset of live leeches was maintained in seawater for transport to the laboratory for immediate microscopic examination. The remaining specimens were dissected into tissue pieces and snap-frozen in liquid nitrogen for subsequent genomic DNA extraction.

### 2.3 Identification of Parasites

For morphological identification, blood smears containing *Trypanosoma* sp. were prepared, air-dried, fixed in methanol, and stained with Giemsa solution (48900-100ML-F, Sigma-Aldrich). The stained smears were examined and photographed under a Nikon Eclipse Ni-U microscope. For molecular analysis, genomic DNA was extracted from distinct parasite sources using standardized protocols. Trypanosoma was isolated from blood samples through a two-step centrifugation process: initial centrifugation at 50 × g for 5 min at 4 °C to remove cellular debris, followed by centrifugation of the supernatant at 1500 × g for 5 min at 4 °C to collect the parasite pellet. DNA extraction from these pellets was performed using the DNeasy Blood & Tissue Kit (69504, Qiagen, Hilden, Germany), while genomic DNA from benedeniid monogeneans and fish leeches was extracted using the Marine Animal Tissue Genomic DNA Extraction Kit (DP324-03, Tiangen, China). All procedures were conducted strictly according to the manufacturers’ protocols. PCR amplification for species identification was carried out using the specific primers listed in Table 1.

**Table 1.**
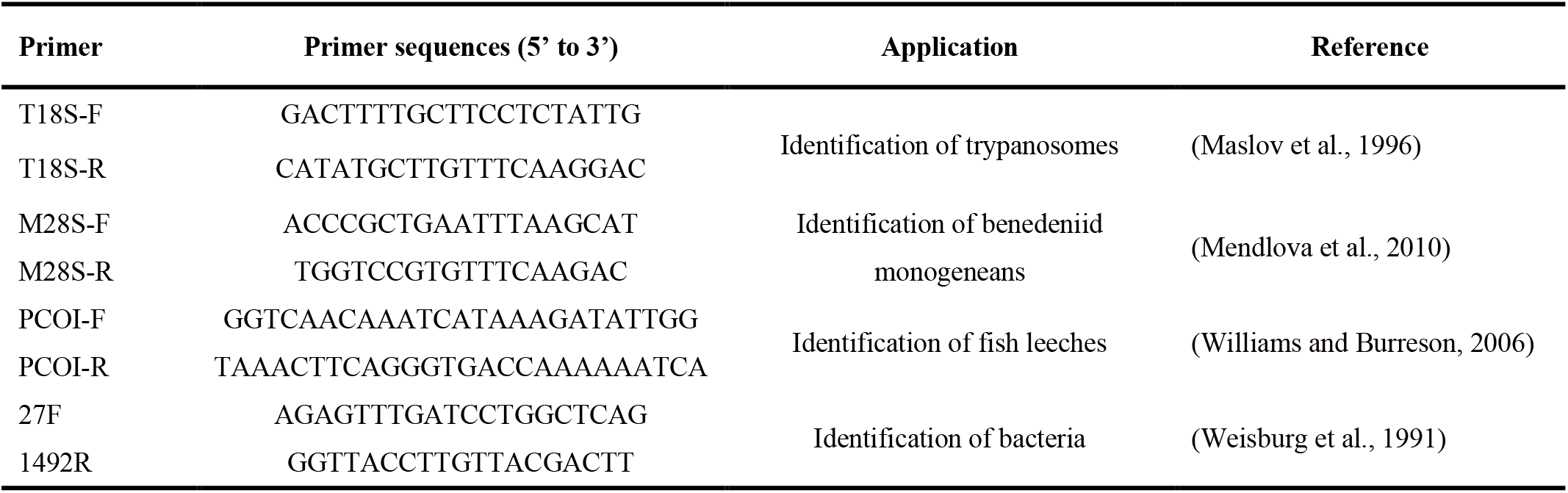
Primers used for pathogen identification in this study.

### 2.4 Isolation and Identification of Pathogenic Bacteria

Following the collection of parasites, diseased fish were dissected in sterile trays to examine and record visceral lesions. Bacterial samples were aseptically collected from the skin, liver, spleen, kidney, intestine, gills, and any necrotic wounds. To minimize surface contamination, surgical instruments (scalpels or scissors) were sterilized by flaming for 3-5 seconds over an alcohol lamp, the broad side of the sterile blade was then pressed against the target tissue surface, followed by incision into the underlying organ. A sterile disposable inoculating loop was inserted into the incised tissue, gently rotated, and subsequently streaked onto Columbia Blood Agar plates (CP0160, HuanKai Microbial, Guangzhou, China). The plates were incubated at 28 °C for 24-48 hours. Single, distinct colonies were selected and transferred to fresh Brain Heart Infusion (BHI) broth, followed by incubation at 28°C in a shaking incubator for 12 hours. Then the PCR amplification was performed using Phanta Max Master Mix (Dye Plus) P525 (Vazyme, Nanjing, China) with universal primers 27F and 1492R (listed in Table 1). Positive PCR products were submitted to Beijing Biomarker Technologies Co., LTD. (Beijing, China) for Sanger sequencing to identify the bacterial species.

Approximately 1 L of water was collected from the aquaculture environment of diseased fish. Microorganisms were concentrated via vacuum filtration using sterile nitrocellulose membranes (0.22 μm pore size, 47 mm diameter) mounted on a multi-channel filtration pump (DCBM-080, DCHENGL, Haining, China). Following filtration, the membranes were aseptically transferred to sterile centrifuge tubes, and environmental DNA was extracted using the TGuide Smart Envir-DNA Kit (DP812, Tiangen, Beijing, China). Concurrently, whole specimens of benedeniid monogeneans and fish leeches were rinsed with sterile water to remove surface contaminants, and total genomic DNA was extracted using the DNeasy Blood & Tissue Kit (69504, Qiagen, Hilden, Germany). DNA samples meeting quality standards were submitted to Beijing Biomarker Technologies Co., LTD. (Beijing, China) for high-throughput microbiome sequencing to identify associated pathogenic bacteria.

### 2.5 Detection of Trypanosomes in benedeniid monogeneans and Fish Leeches

Live specimens of benedeniid monogeneans and fish leeches were dissected under an Olympus SZX16 stereomicroscope. Intestinal contents were smeared onto glass slides and examined for the presence of Trypanosomes using a Nikon Eclipse Ni-U microscope. For molecular detection, total genomic DNA was extracted from whole individuals of benedeniid monogeneans and fish leeches using the DNeasy Blood & Tissue Kit (69504, Qiagen, Hilden, Germany), strictly following the manufacturer’s instructions. PCR amplification was subsequently performed using Trypanosome-specific primers T18S-F/R (Table 1) to confirm the presence of hemoflagellates within these ectoparasites. PCR was performed in 25 μL mixtures containing 1μL of DNA, 0.5 μL of each primer pair, 18 μL of ddH_2_O, 2 μL of dNTP Mixture, 0.5 μL of Ex-Taq enzyme (RR001B, Takara, Dalian, China) and 2.5 μL Buffer. The PCR program consisted of one cycle at 95 ℃ for 30s, followed by 40 cycles of amplification at 95 ℃ for 5 s, 60 ℃ for 30s and 72 ℃ for 1 min, and finally at 72 ℃ for 10 min. To check the integrity of the PCR products, the products were electrophoresed on 1.0% agarose gel and viewed by a Molecular Imager Gel Doc XR system (Bio-Rad, USA).

### 2.6 Experimental Simulation of *T. carassii spectrum* Transmission by leech

*M. salmoides* (body length 7.38 ± 0.42 cm, weight 4.48 ± 0.57 g) were purchased from Hengxing (Guangzhou) Fisheries Development Co., Ltd., Yangchun Branch (Yangchun, Guangdong, China). The fish were monitored in indoor laboratory fish tanks with recirculating fresh water at 25 ℃ for a month prior to infection experiments, with blood samples screened via microscopy to ensure the absence of haemoflagellate infections. The *T. carassii spectrum* strain used in this study was previously isolated and maintained in our laboratory (Wang et al., 2025). As illustrated in Figure 1, the transmission simulation experiment consisted of three groups (A, B, and C), each containing 15 healthy *M. salmoides*. Initially, 15 fish per group were intraperitoneally injected with *T. carassii spectrum* and housed in 10 L glass tanks (35 cm × 20 cm × 23 cm). The experiment commenced when the parasitemia in these infected fish reached 10^8^/mL. At this point, 15 additional healthy, marked *M. salmoides* were introduced into each tank for co-culture. The experimental conditions varied as follows: Group A involved the placement of one *Poecilobdella manillensis* (a blood-feeding leech vector) on each infected fish; Group B served as a co-culture control without leeches; and Group C served as a physical separation control, where infected and healthy fish were separated by plastic partitions to prevent direct contact, also without leeches (Figure 1). During the 14-day co-culture period, feeding was reduced by half, and siphoning was performed every two days to maintain water quality. Upon completion of the experiment, blood samples were collected from the healthy fish in all three groups. Infection status was assessed through blood smear microscopy and PCR analysis.

**Figure 1.**
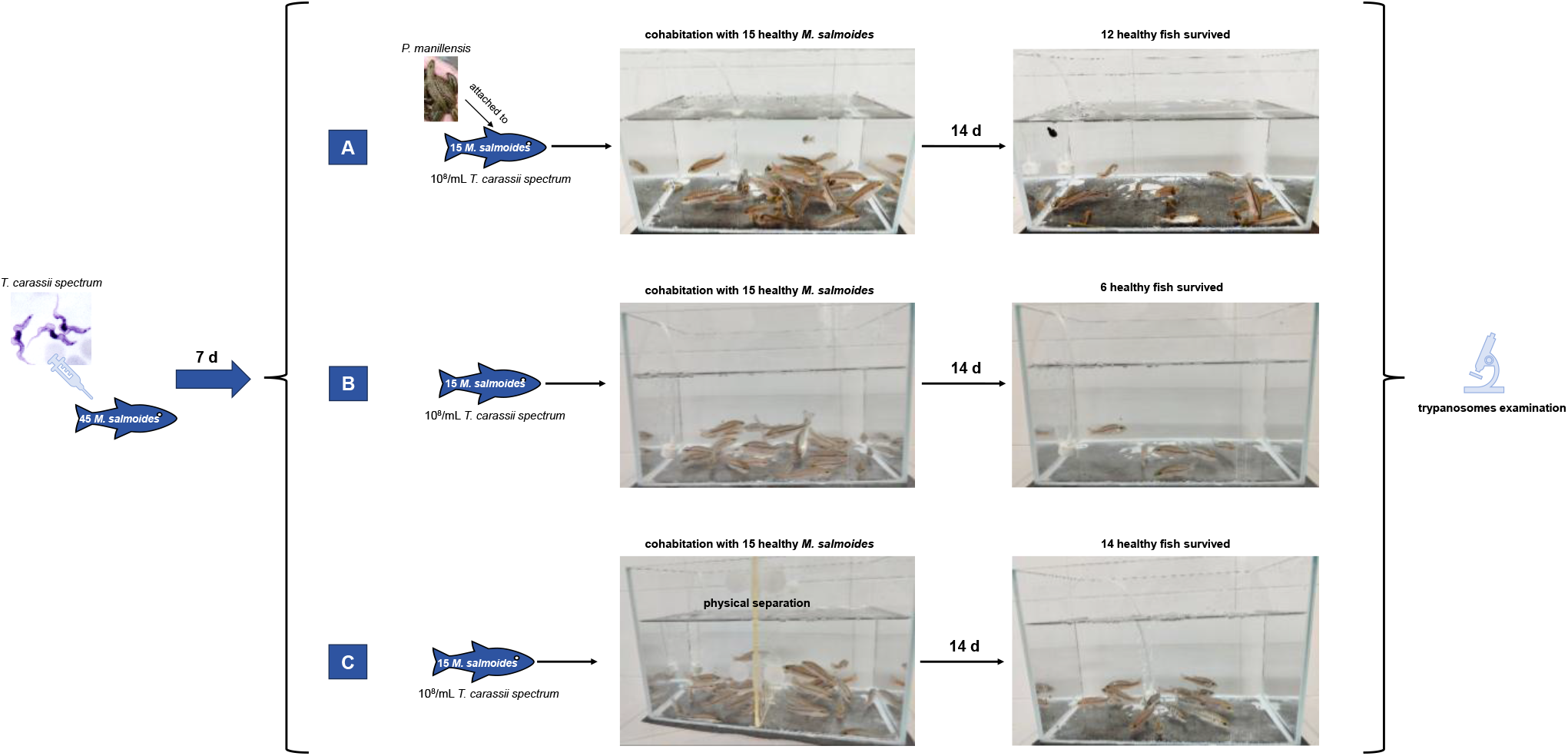
Schematic diagram of the experimental simulation of *Trypanosoma carassii spectrum* transmission mediated by leeches.

### 2.7 In vitro Feeding Assay of *N. girellae* with *T. carassii spectrum*

Adult *N. girellae* were gently detached from infected host fish using sterile plastic forceps. Each worm was immediately transferred to a glass slide pre-loaded with a single drop of sterile seawater to maintain osmotic balance. Subsequently, 10 μL of blood containing *T. carassii spectrum* (approximately 10^8^/mL) was applied to the surface of the worm. The feeding behavior and potential uptake of trypanosomes were observed in real-time and documented using an Olympus SZX16 dissecting microscope equipped with a digital camera.

## 3. Results

### 3.1 Field survey

As is shown in Figure 2, the outbreak occurred in Ningde waters, Fujian Province, during a period of elevated water temperature (29°C). Affected *L. crocea* (body length: 14.87 ± 1.72 cm) displayed severe clinical signs, including extensive epidermal erosion,’crater-like’ ulcerative lesions, ocular opacity or hyperemia, and fin rot (Fig. 2C). Behavioral observations noted lethargy and frequent rubbing against net cages, indicative of ectoparasitic irritation. The reported mortality rate exceeded 50%, with numerous dead fish floating on the water surface (Fig. 2B). Parasitological examination of five randomly selected diseased fish revealed a 100% co-infection rate with *Neobenedenia* sp. and *Trypanosoma* sp. (Table 2). The mean infection intensity of *Neobenedenia* sp. was 16.40 ± 10.13, while trypanosomemia in peripheral blood exceeded 10^8^/mL (Fig. 3). Bacterial isolation from tissue samples identified *Bacillus cereus* and *Photobacterium damselae* as the predominant pathogens, with a prevalence of 80% for each species. Additionally, *Vibrio cholerae* and *Vibrio harveyi* were isolated from affected tissues (Table 2; Fig. S1; Supplementary file 1), suggesting a complex polyparasitic and bacterial etiology associated with the mortality event.

**Figure 2.**
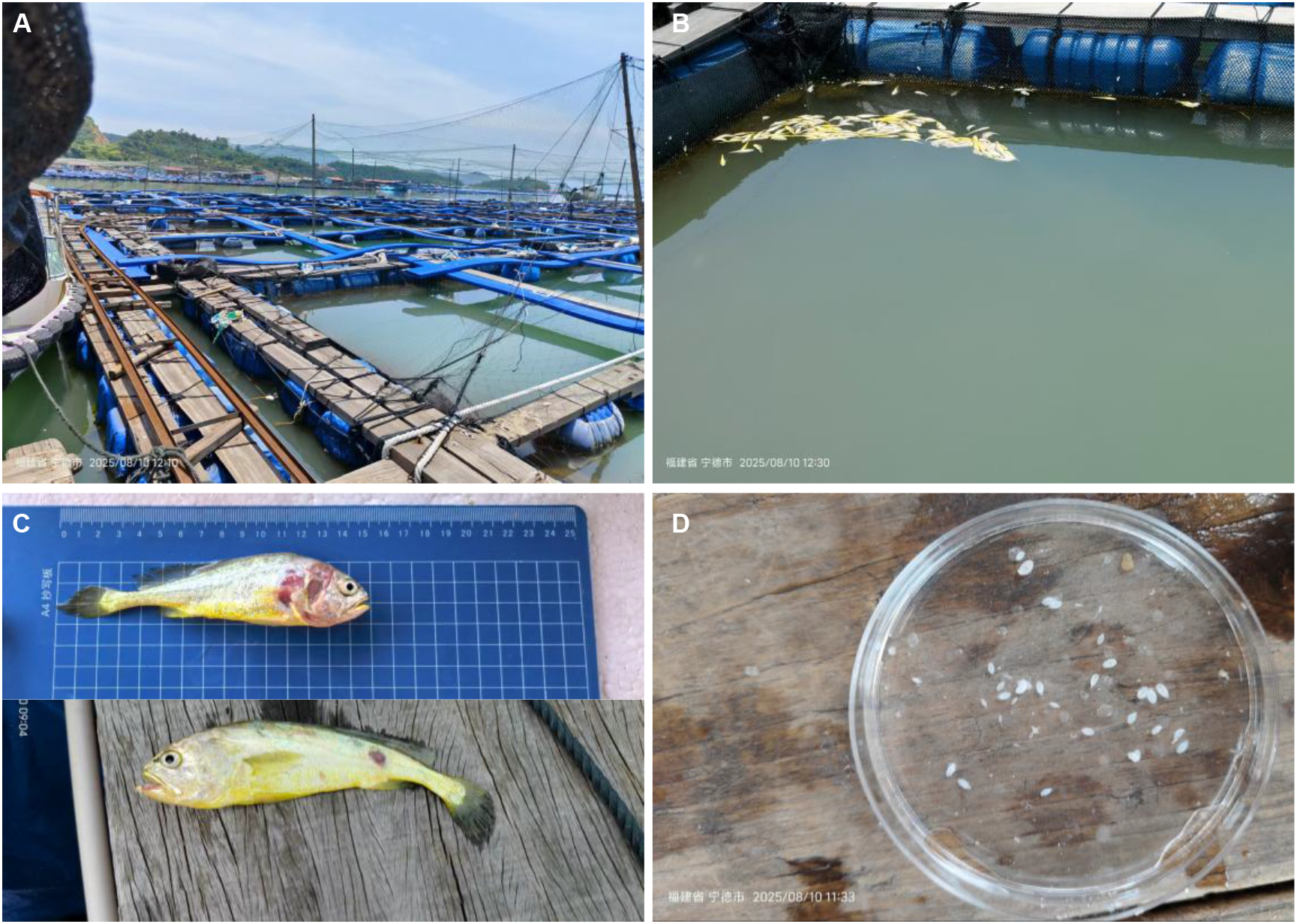
Overview of the sampling location and pathological findings in Ningde, Fujian. (A) Floating net cages used for *Larimichthys crocea* aquaculture; (B-C) Representative images of diseased and moribund *L. crocea* exhibiting clinical signs of infection; (D) *Neobenedenia girellae* parasites collected from infected hosts.

**Figure 3.**
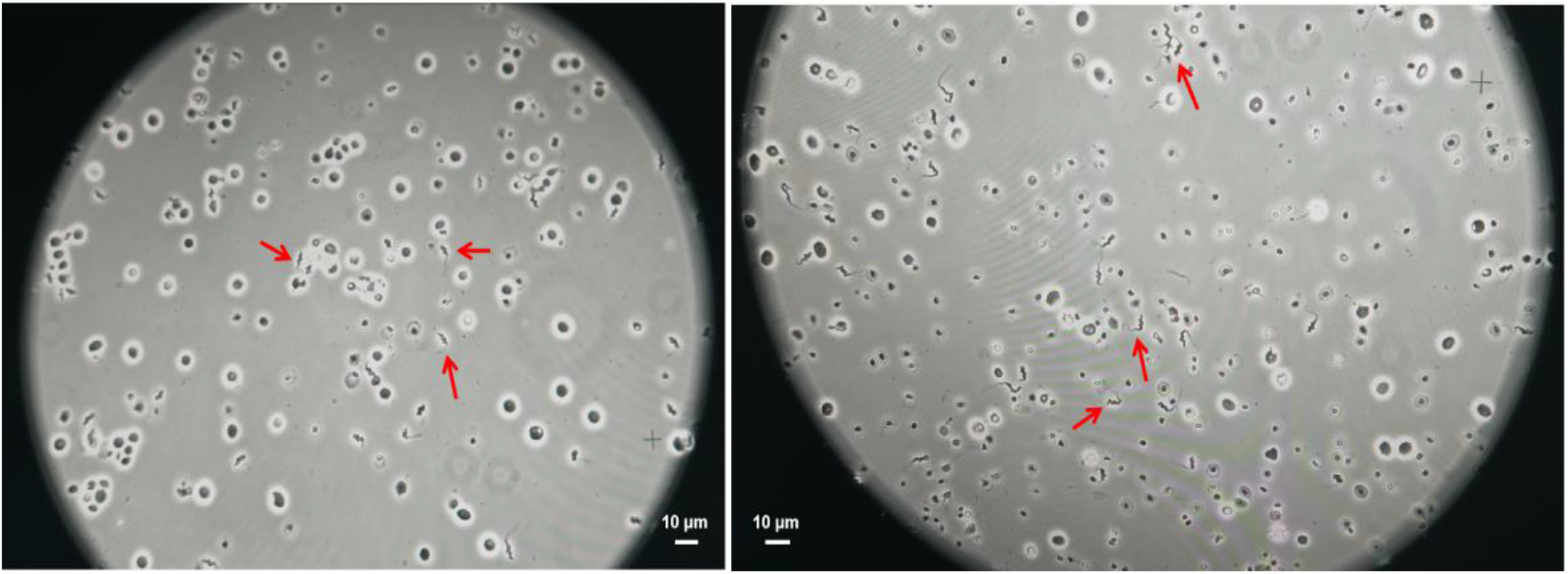
Microscopic examination of blood smears from *Larimichthys crocea* collected in Ningde, Fujian Province. Red arrows indicate *Trypanosoma carassii spectrum*. Scale bars = 10 μm.

**Table 2.**
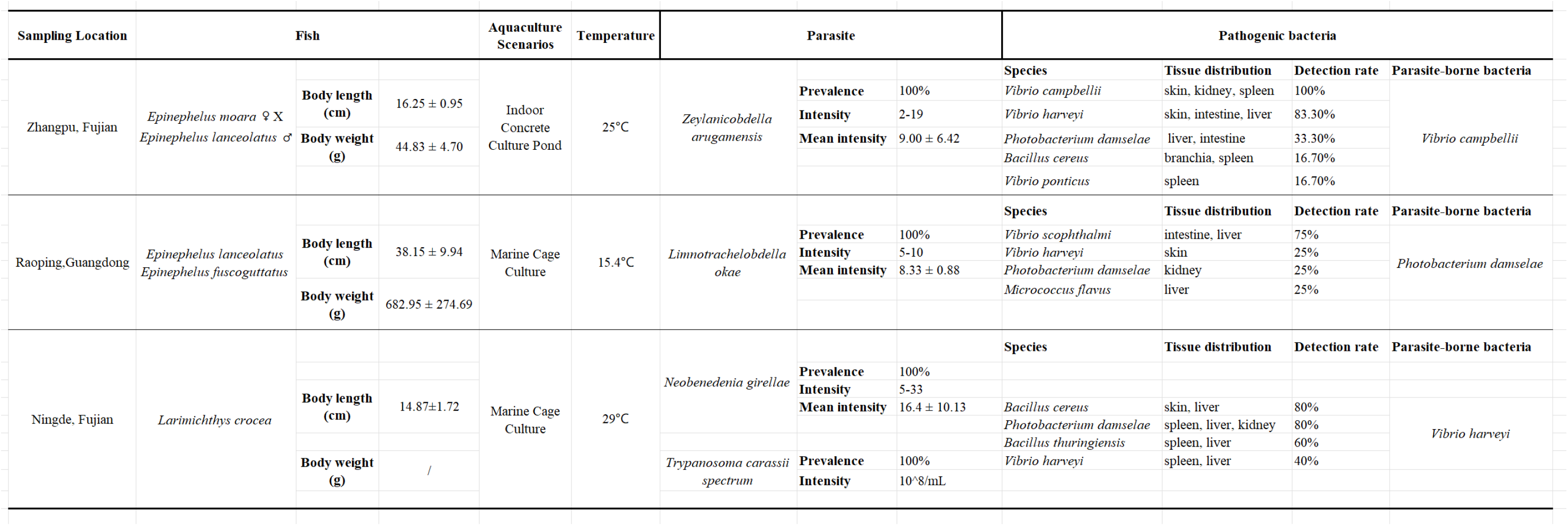
Identification of pathogens from host and vector samples collected during three field surveys in Fujian and Guangdong provinces.

The disease outbreak occurred in Zhangpu, Fujian Province, affecting hybrid grouper (*Epinephelus moara* ♀ X *Epinephelus lanceolatus* ♂) with a mean body length of 16.25 ± 0.95 cm and body weight of 44.83 ± 4.70 g at a water temperature of 25°C. Clinically affected fish exhibited lethargy, moribund states, and abdominal dropsy, with numerous individuals floating belly-up on the water surface (Fig. 4A). Macroscopic examination revealed aggregations of piscine leeches (approximately 2 cm in length) attached to scaleless regions, such as the fin bases and ventral abdomen, often displaying inchworm-like locomotion (Fig. 4B, C). Parasitological analysis of six sampled fish demonstrated a 100% prevalence of leech infection, with a mean intensity of 9.00 ± 6.42 (Table 2). Bacterial isolation from diseased tissues identified *Vibrio campbellii* and *Vibrio harveyi* as the predominant pathogens, each with a prevalence exceeding 80%. Additionally, *Photobacterium damselae*, *Bacillus cereus*, and *Vibrio ponticus* were isolated (Table 2; Fig. S2; Supplementary File 2).

**Figure 4.**
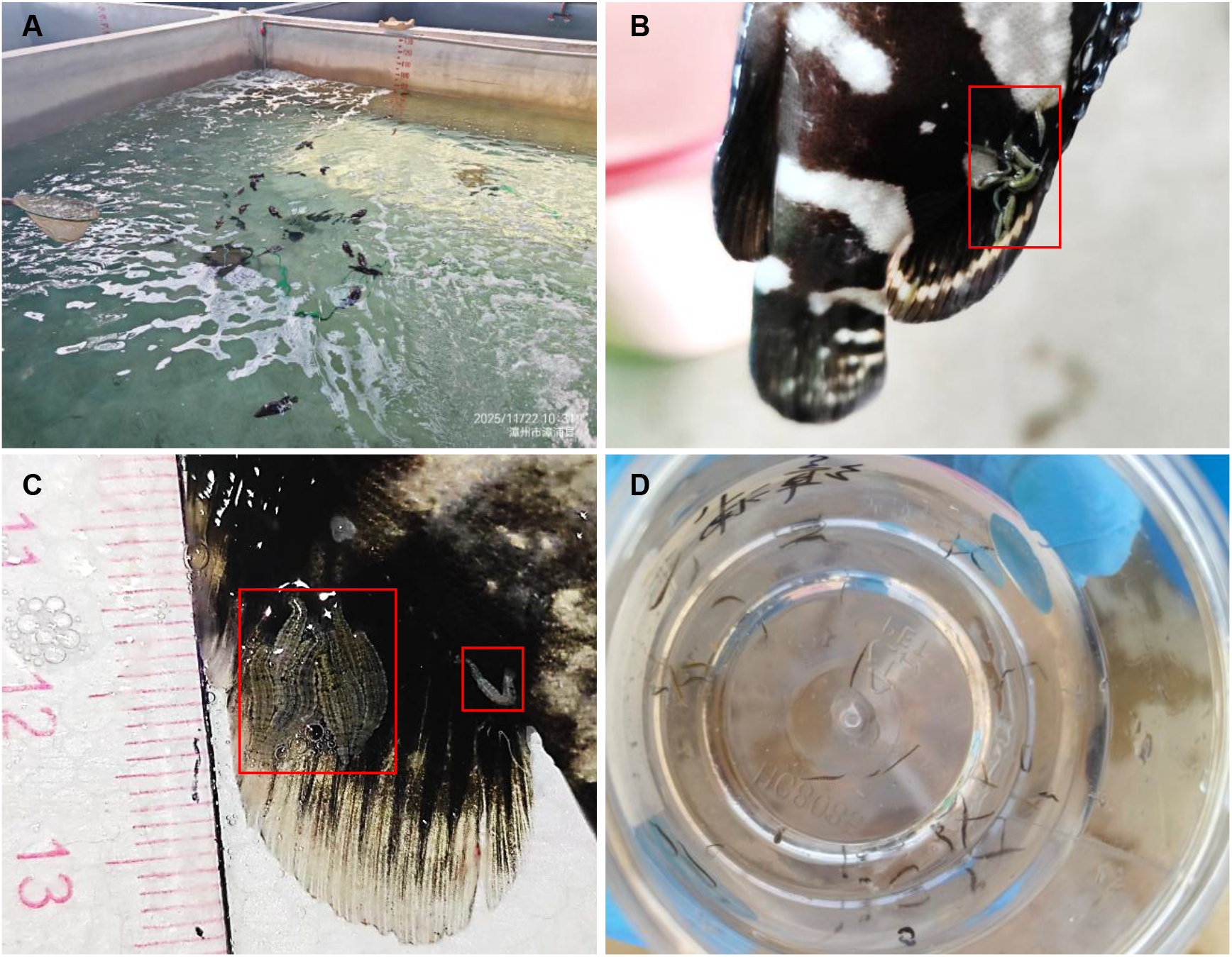
Sampling context in Zhangpu, Fujian Province. (A) Aquaculture pond with clinically affected groupers. (B-C) Aggregations of *Zeylanicobdella arugamensis* on the dorsal fins of infected groupers; red boxes highlight individuals of *Z. arugamensis*. (D) Collected *Z. arugamensis* specimens.

The disease outbreak in Raoping, Guangdong Province, primarily affected grouper species, including the *Epinephelus lanceolatus* and *Epinephelus fuscoguttatus* (Fig. 5A). The infected fish had a mean body length of 38.15 ± 9.94 cm and a mean body weight of 682.95 ± 274.69 g, with the outbreak occurring at a water temperature of 15°C. Macroscopic examination revealed piscine leeches attached to scaleless or sparsely scaled regions, such as the fin bases, posterior margin of the operculum, and ventral abdomen (Fig. 5B, C). Juvenile leeches measured approximately 2-5 cm in length, while adults exceeded 10 cm. Affected fish exhibited multiple open wounds on the body surface, with some showing hemorrhage or ulceration; juvenile leeches were observed emerging from larger lesions (Fig. 5C). Parasitological analysis of eight sampled fish demonstrated a 100% prevalence of leech infection, with a mean intensity of 8.33 ± 0.88 individuals per host (Table 2). Bacterial isolation from diseased tissues identified *Vibrio scophthalmi* as the predominant pathogen, with a prevalence of 75%. Additionally, *Vibrio harveyi*, *Photobacterium damselae*, and *Micrococcus flavus* were isolated from the affected tissues (Table 2; Fig. S3; Supplementary File 3).

**Figure 5.**
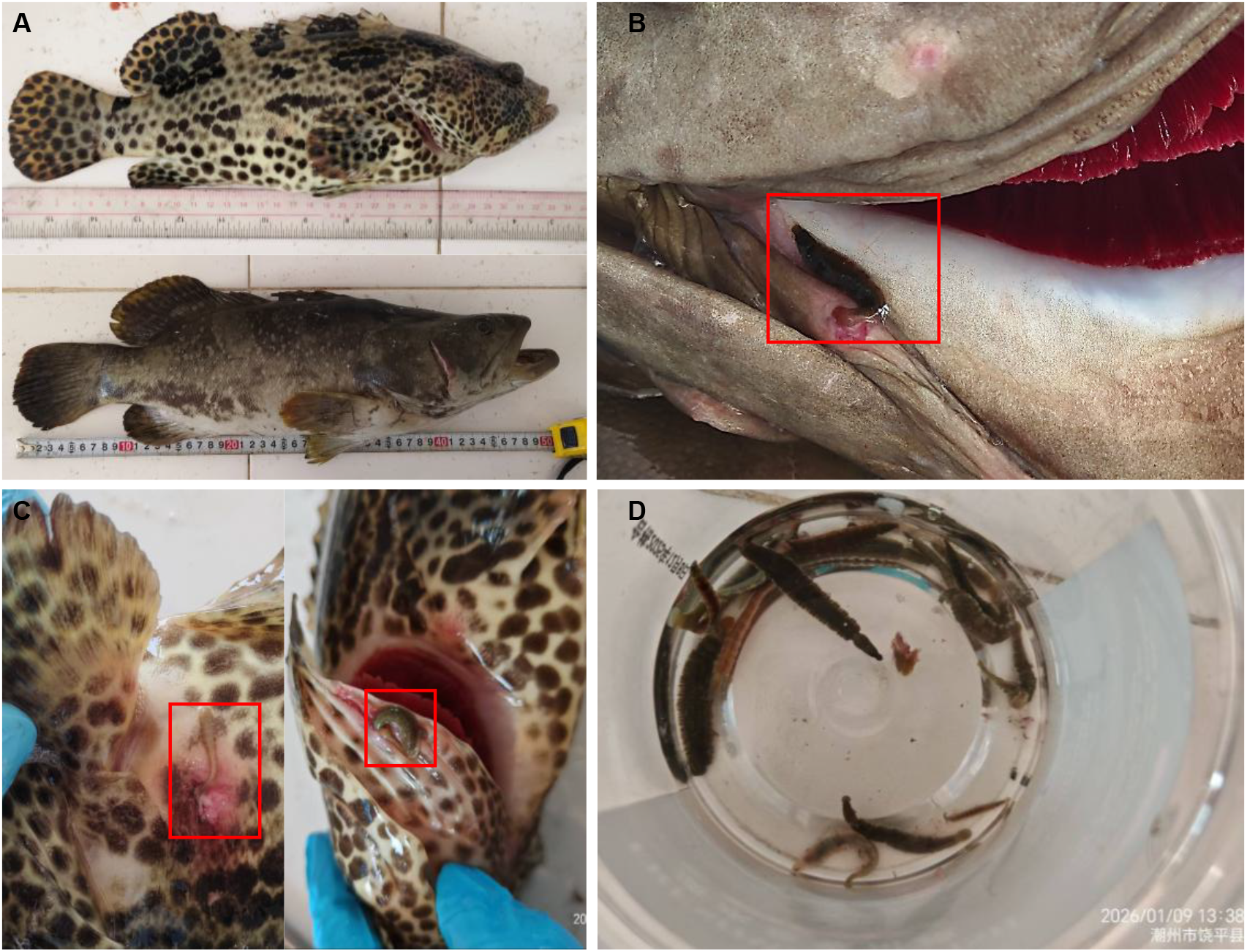
Sampling context in Raoping, Guangdong Province. (A) Clinically affected groupers exhibiting disease symptoms. (B-C) Aggregations of *Limnotrachelobdella okae* on infected groupers; red boxes highlight individual leeches. (D) Collected *L. okae* specimens.

### 3.2 Vector Potential of Parasites Identified in Field Surveys

Molecular characterization identified the trypanosomes isolated from diseased *L. crocea* in Ningde, as belonging to the *Trypanosoma carassii spectrum* (Fig. 6D; Supplementary File 1), while the *Neobenedenia* sp. was identified as *Neobenedenia girellae* (Fig. 6C; Supplementary File 1). Notably, *Vibrio harveyi*, the predominant pathogen isolated from diseased fish, was also detected within these parasites (Table S1). Intriguingly, molecular traces of *T. carassii spectrum* were detected in *N. girellae* specimens collected from two individual hosts (Fig. 6E; Supplementary File 1). *In vitro* observations confirmed that *N. girellae* ingests both *T. carassii spectrum* along with host erythrocytes (Video S1), maintaining trypanosome viability for several hours post-ingestion (Video S2). Fish leeches collected from diseased groupers in Zhangpu were identified as *Zeylanicobdella arugamensis* (Fig. 6A; Supplementary File 2), whereas those from Raoping were identified as *Limnotrachelobdella okae* (Fig. 6B; Supplementary File 3). Extensive microscopic and molecular screening revealed no evidence of trypanosome infection in either leech species. However, molecular analysis revealed the presence of *Vibrio campbellii* in *Z. arugamensis* (Table S2) and *Photobacterium damselae* in *L. okae* (Table S3), respectively, mirroring the bacterial pathogens isolated from the corresponding diseased host tissues. Control water samples collected during the three sampling events tested negative for these specific pathogens.

**Figure 6.**
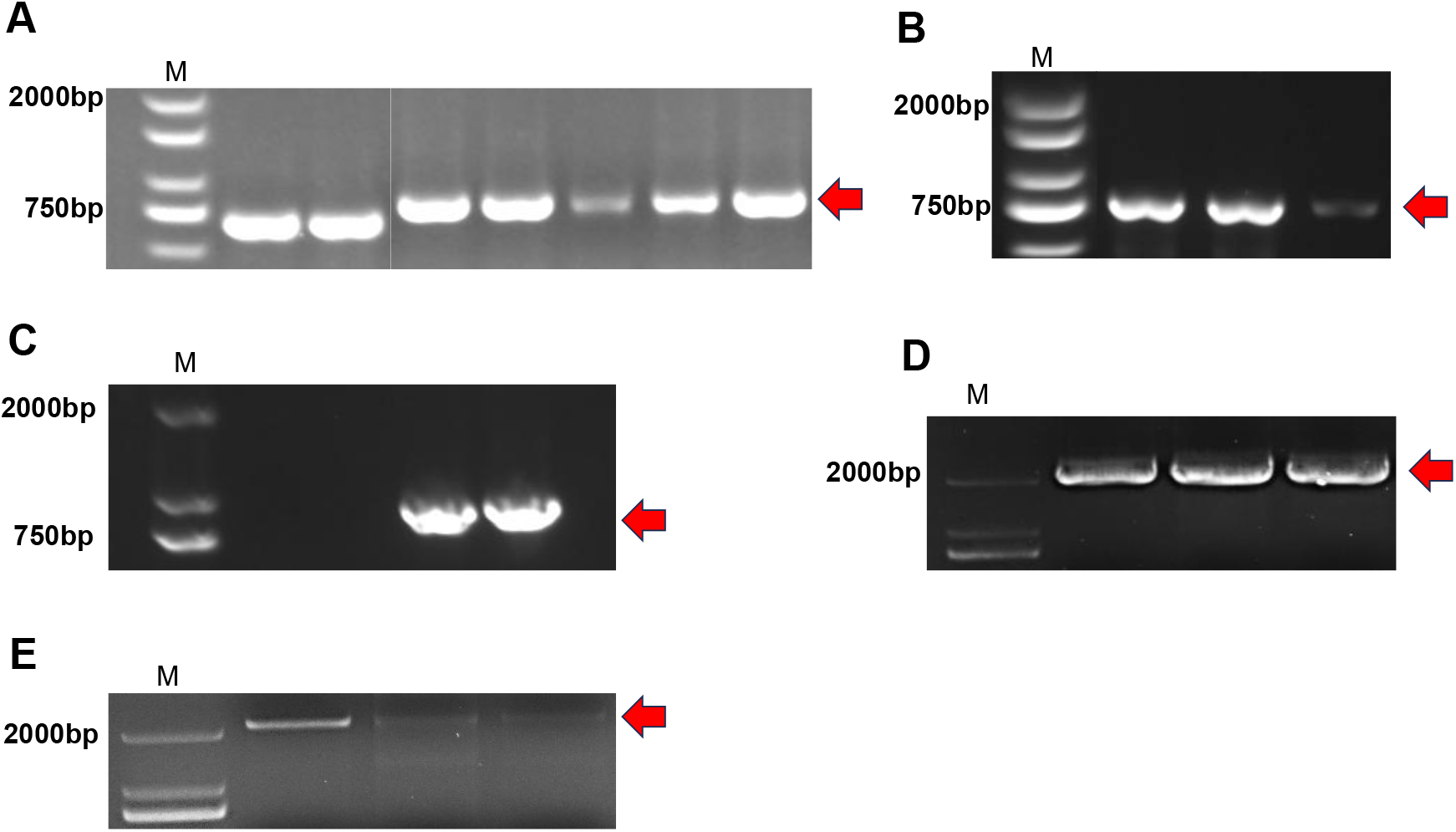
Agarose gel electrophoresis results of PCR amplification for parasite identification. M: DNA marker. Red arrows indicate the target bands. (A) Identification of fish leeches collected from Zhangpu, targeting the COI gene. (B) Identification of fish leeches collected from Raoping, targeting the COI gene. (C) Identification of *Neobenedenia* sp. collected from Ningde, targeting the 28S rRNA gene. (D) Identification of trypanosomes collected from Ningde, targeting the 18S rRNA gene. (E) Detection of trypanosome traces within *Neobenedenia* sp. collected from Ningde, Fujian, targeting the 18S rRNA gene.

3.3 Experimental Evidence for Leech-Mediated Transmission of *T. carassii spectrum*

All initial carrier *M. salmoides* died within 3 days post-cohabitation. In Group A, all *P. manillensis* migrated to the healthy fish. During the experiment, six leeches were consumed by the fish. Three healthy fish died unexpectedly during the early phase of the experiment, with no trypanosome infection detected in any of them. At the conclusion of the study, 12 healthy fish in Group A remained. PCR analysis revealed that 7 out of 12 healthy bass in Group A were positive for *T. carassii spectrum* (Fig. 7B), yielding an infection rate of 58.33%. Blood smears confirmed trypanosome presence in 6 of these 7 individuals (excluding A10 due to low parasitemia), with parasite densities exceeding 10^7^/mL (Fig. 7A). In Group B, 6 out of 15 healthy fish tested PCR-positive (Fig. 8B), resulting in a 40.00% infection rate; all positive cases showed detectable trypanosomes in blood smears with densities above 10^7^/mL. In contrast, no trypanosome infections were detected in Group C (Fig. 9). These findings demonstrate that *T. carassii spectrum* can be transmitted via leech vectors and potentially through direct host contact, but not through waterborne transmission in the absence of contact. Notably, leech-mediated transmission exhibited higher efficiency.

**Figure 7.**
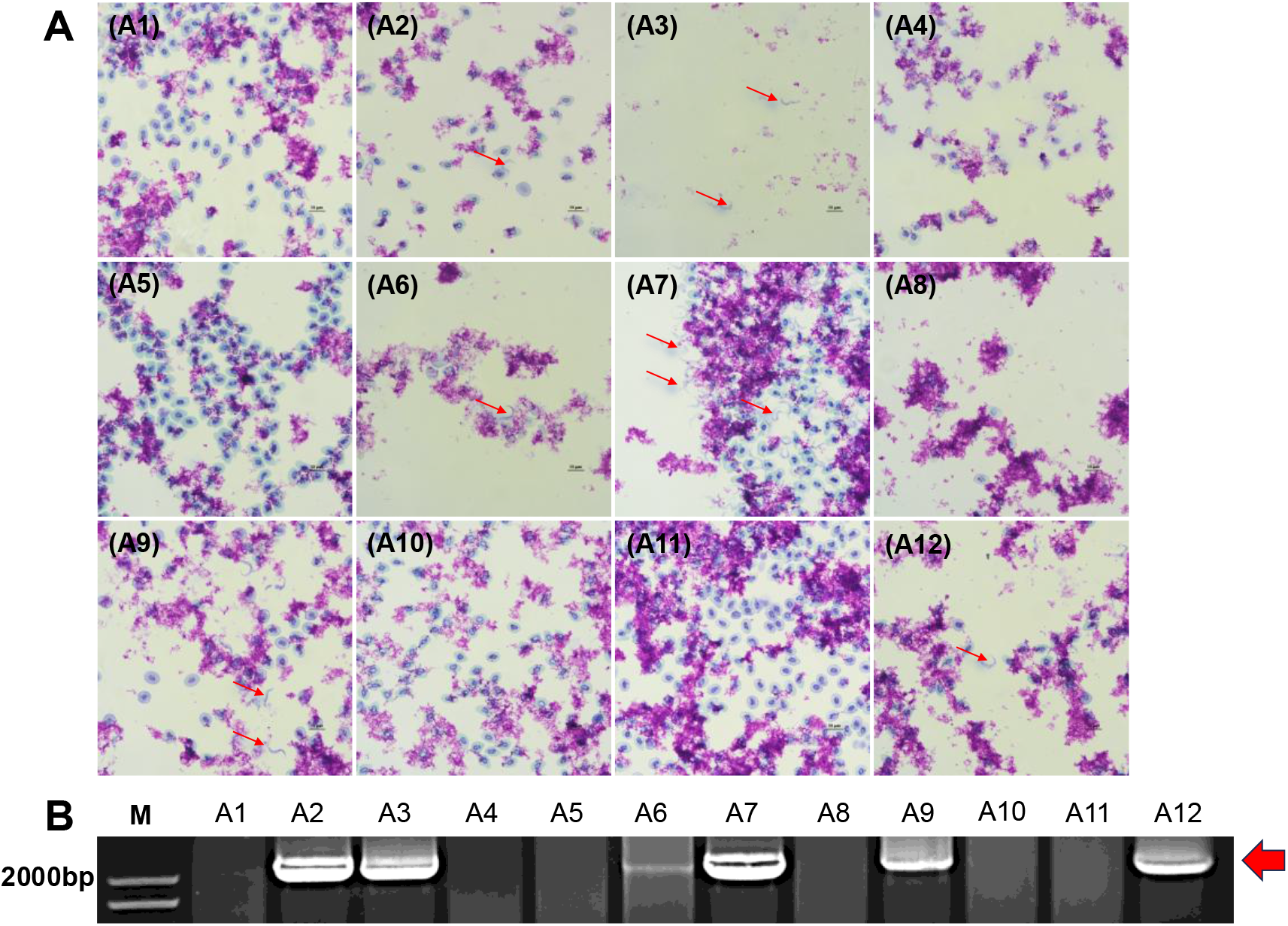
Detection of *T. carassii spectrum* infection in healthy *M. salmoides* from Group A. (A) Photomicrographs of Giemsa-stained blood smears showing trypanosome morphology; red arrows indicate *T. carassii spectrum*; Scale bars = 10 μm. (B) Agarose gel electrophoresis results of PCR amplification targeting the 18S rRNA gene from fish blood samples; M: DNA marker; red arrow indicates the specific amplicons corresponding to *T. carassii spectrum* DNA.

**Figure 8.**
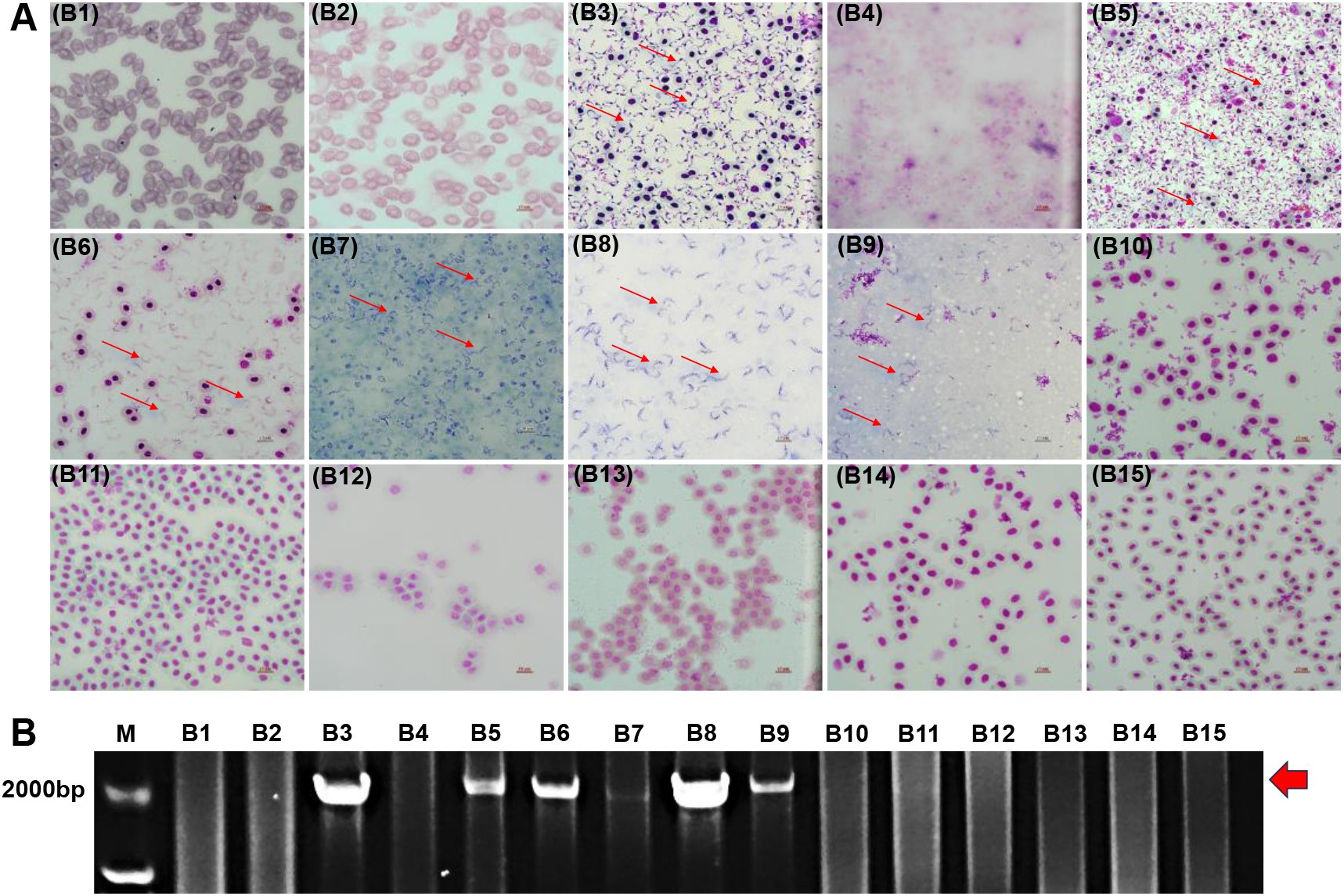
Detection of *T. carassii spectrum* infection in healthy *M. salmoides* from Group B. (A) Photomicrographs of Giemsa-stained blood smears showing trypanosome morphology; red arrows indicate *T. carassii spectrum*; Scale bars = 10 μm. (B) Agarose gel electrophoresis results of PCR amplification targeting the 18S rRNA gene from fish blood samples; M: DNA marker; red arrow indicates the specific amplicons corresponding to *T. carassii spectrum* DNA.

**Figure 9.**
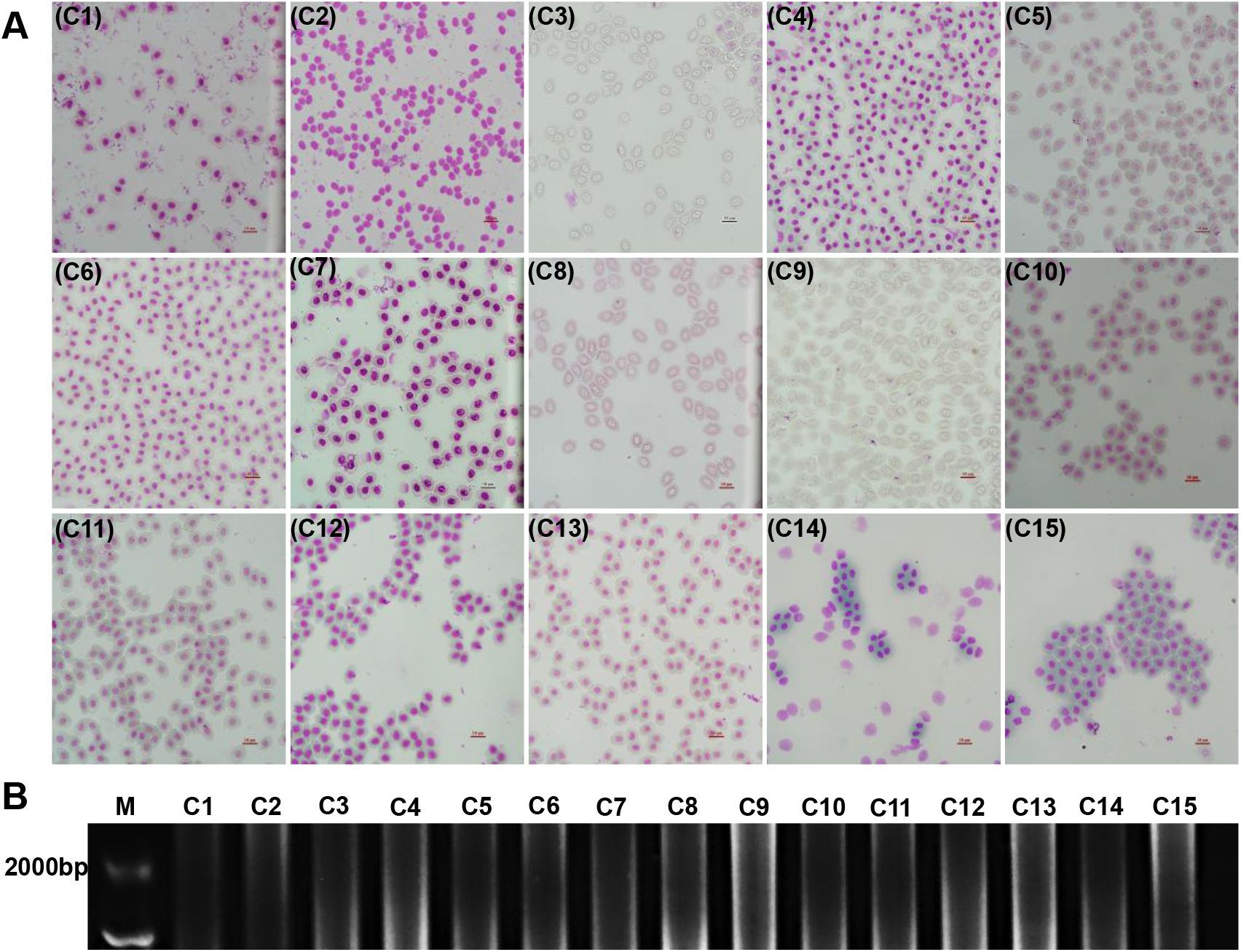
Detection of *T. carassii spectrum* infection in healthy *M. salmoides* from Group C. (A) Photomicrographs of Giemsa-stained blood smears showing trypanosome morphology; Scale bars = 10 μm. (B) Agarose gel electrophoresis results of PCR amplification targeting the 18S rRNA gene from fish blood samples;M: DNA marker.

## 4. Discussion

In aquaculture systems, parasites not only act as direct pathogens causing tissue damage and growth impairment but also serve as critical biological vectors or synergistic factors, significantly exacerbating the transmission and pathogenicity of viral, bacterial and other microbial agents (Islam et al., 2024; Okon et al., 2023). This’collateral disease’ effect is often overlooked by traditional single-pathogen perspectives, leading to ineffective disease control strategies (Bouwmeester et al., 2021). Previous studies have demonstrated that parasites serve as vectors for various pathogens, functioning either as mechanical vectors - wherein pathogens are merely transported without biological development, such as monogeneans carrying agents on their surfaces or within their bodies (Cardoso et al., 2022); or as biological vectors, where pathogens undergo development or proliferation, exemplified by trypanosomes in leeches (Smit et al., 2020); In certain instances, specific pathogens rely exclusively on these parasitic vectors for transmission (Serem et al., 2024; Strand and Burke, 2014).

Since the latter half of 2023, a novel subspecies of *Trypanosoma carassii*, characterized by its broad host range (*Trypanosoma carassii spectrum*), has caused severe trypanosomiasis outbreaks in cage-cultured *L. crocea* in Sandu Bay, Ningde, Fujian Province. This epidemic has rapidly spread along the southeastern coast of China, resulting in substantial economic losses to the aquaculture industry. Notably, this represents the first recorded instance of trypanosomiasis in *L. crocea*, with phylogenetic evidence suggesting a freshwater origin for the pathogen (Liu et al., 2026). Despite extensive investigations, the source and transmission routes of *T. carassii spectrum* remain elusive. Trypanosomes are obligate blood parasites of vertebrates that lack the capacity for active host exit, relying predominantly on hematophagous invertebrates (particularly leeches) for transmission (Krige et al., 2019). In the established life cycle of *T. carassii*, the leech *Hemiclepsis marginata* serves as the primary vector, where trypomastigotes transform into short, stout forms, differentiate into epimastigotes, and ultimately develop into infectious metacyclic forms within the leech’s proboscis sheath (Lom and Dyková, 1992). However, extensive surveys since the onset of the *T. carassii spectrum* outbreak have failed to detect any leech populations in either infected hosts or the surrounding aquaculture environment, leaving the actual transmission route in this marine system unresolved.

In this study, *Z. arugamensis* and *L. okae* leeches were collected from waters adjacent to the *L. crocea* trypanosomiasis outbreak zones. *Zeylanobdella arugamensis* is predominantly distributed across Southeast Asia and the southeastern coastal waters of China, exhibiting an optimal thermal preference of approximately 26°C. In Chinese aquaculture systems, this leech species primarily parasitizes groupers (*Epinephelus* spp.) and has been implicated in disease outbreaks in Hainan Province (Wang et al., 2018). Notably, *Z. arugamensis* has been documented as an intermediate host for the marine fish trypanosome *Trypanosoma nudigobii*, establishing its vectorial competence for trypanosome transmission in marine environments (Hayes et al., 2014). Historically, the *L. okae* was predominantly distributed in northern waters, including the Yellow Sea of China, Tokyo Bay in Japan, and Peter the Great Bay in Russia (Yang, 1996). This leech infects a variety of marine fish hosts, such as *Miichthys miiuy*, *Epinephelus* spp. and *Seriola dumerili*. Characterized by euryhaline tolerance, *L. okae* can survive in both freshwater and seawater, with an optimal water temperature range of 5-15°C. Notably, around 2023, disease outbreaks associated with *L. okae* were reported for the first time in the southern coastal waters of China (Che et al., 2026). If *L. okae* migrated southward from its northern habitats driven by ocean currents and other environmental factors, its migration route would spatially align with the epidemic zones of *T. carassii spectrum* in *L. crocea*. Regrettably, our investigation failed to reveal any traces of trypanosomes within the two leech species surveyed, this issue may require investigation with a larger sample size for resolution. Furthermore, particular attention should be paid to these fish leeches. Beyond the direct harm they inflict on marine-cultured fish, vigilance is crucial to prevent the transmission of *T. carassii spectrum* by these vectors to a broader range of marine fish hosts and into wider marine environments.

Following the initial isolation of *T. carassii spectrum* from *L. crocea*, our team attempted to establish an indoor infection model to facilitate long-term research. However, this effort was hindered by the inability to maintain *L. crocea* in small-scale indoor systems for extended periods. Furthermore, *T. carassii spectrum* exhibited poor survival in other marine hosts amenable to laboratory recirculating aquaculture systems, such as *Epinephelus awoara* and *Acanthopagrus latus*, thereby precluding the development of a marine fish-based experimental model. Fortunately, we identified that *T. carassii spectrum* is capable of infecting various freshwater species, including *M. salmoides*, *Oreochromis niloticus* and koi carp, with *M. salmoides* demonstrating the highest susceptibility (Wang et al., 2025). Concurrently, the freshwater leech *P. manillensis* was observed to attach to and survive on largemouth bass. Consequently, we established a transmission model comprising *T. carassii spectrum*, *M. salmoides* and *P. manillensis*. Our results indicate that this model successfully recapitulates the complete transmission cycle of *T. carassii spectrum*, with *P. manillensis* exhibiting high transmission efficiency. This elevated efficiency may be attributed to the non-parasitic, transient nature of *P. manillensis*, which facilitates frequent transfer among co-housed largemouth bass (Enguang, 2008). Additionally, the aggressive behavior of largemouth bass, characterized by frequent biting and attacking of leeches on the fish surface, likely contributes to mechanical dissemination. Although we have observed that *T. carassii spectrum* can survive within *P. manillensis* for more than 24 hours, but systematic quantitative data regarding this duration remain lacking. Current findings confirm that leeches could serve as mechanical vectors for the carriage and transmission of *T. carassii spectrum*. However, further field investigations and systematic experimental studies are required to elucidate the specific transmission pathways of *T. carassii spectrum* in marine environments.

In this study, we unexpectedly identified that *N. girellae* and direct host-to-host contact represent potential transmission routes for *T. carassii spectrum*. In high-density aquaculture systems, physical interactions such as rubbing or mutual biting among host fish, may induce open wounds or subclinical hemorrhages on the fish integument, thereby facilitating the egress of trypanosomes (Bu et al., 2026). It is hypothesized that susceptible hosts may subsequently become infected through the ingestion of free-swimming trypanosomes that exhibit transient survival in the water column. *Neobenedenia girellae*, which inhabit the epidermis of marine fishes, primarily feed on host epithelial cells and mucus, while also ingesting small amounts of blood (AMIIN et al., 2024; LA Tubbs et al., 2005; Whittington, 2011). This hematophagous behavior creates opportunities for the parasite to encounter and mechanically carry trypanosomes. However, given the limited horizontal dispersal capability of *N. girellae*, characterized by strong attachment to the host and rare detachment except upon host mortality (Valles-Vega et al., 2019), we speculate that *N. girellae* serves merely as an incidental vector for *T. carassii spectrum* transmission. Consequently, more robust evidence chains are required to definitively validate these two potential transmission pathways.

In addition, pathogenic bacteria identical to those isolated from diseased fish were detected in the collected *N. girellae* specimens and two species of fish leeches. Accumulating evidence indicates that parasites, especially macroparasites, serve as vectors for the carriage and transmission of secondary pathogens within aquaculture systems. Studies have demonstrated that *Myzobdella lugubris* is a carrier of infectious Viral Hemorrhagic Septicemia virus (VHSV) genotype IVb, positioning it as a potential mechanical vector and environmental reservoir in the Great Lakes ecosystem (Faisal and Schulz, 2009). salmon louse, *Lepeophtheirus salmonis* serves as a vector for *Aeromonas salmonicida* subsp. *salmonicida*, acquiring the pathogen via diverse mechanisms and demonstrating significant transmission potential (Novak et al., 2016). Monogenean *Gyrodactylus cichlidarum* parasitizing tilapia possess a resilient microbial community independent of the host and ambient water, enabling the persistent carriage of *Aeromonas* spp. a and implicating them as the principal vectors for its transmission (Garcia-Vásquez et al., 2026). Consequently, parasites warrant increased attention in aquaculture health management. They can harbor pathogens and coexist with hosts for prolonged periods, leading to sudden disease outbreaks under favorable conditions. Therefore, rigorous monitoring and control of parasites during cultivation may significantly enhance the overall efficacy of disease prevention and control strategies.

## 5. Conclusions

Field surveys and laboratory transmission assays demonstrated that the *T. carassii spectrum* can be transmitted via leech vectors and potentially through direct contact among host fish, with leech-mediated transmission exhibiting higher efficiency. While *N. girellae* can harbor *T. carassii spectrum* internally, this suggests a potential incidental transmission route, particularly in leech-free disease outbreak areas. However, the failure to fully reproduce *N. girellae*-mediated transmission under current experimental conditions, and no leech vectors associated with *T. carassii spectrum* have been identified in disease-endemic waters highlight the need for further research to definitively elucidate the transmission pathways of *T. carassii spectrum*.

Both *N. girellae* and fish leeches harbor fish pathogens, including *V. harveyi* and *P. damselae*, suggesting they may serve as primary vectors for these bacteria. Beyond their direct pathogenicity, parasites may act as vehicles for secondary pathogens, potentially driving disease outbreaks in high-density intensive aquaculture systems. Therefore, integrated parasite control is crucial for enhancing overall fish health management efficiency.

## Supplementary Materials

Figure S1: Streak plate isolation of pathogenic bacteria from various tissues of diseased fish in Ningde, Fujian; Figure S2: Streak plate isolation of pathogenic bacteria from various tissues of diseased fish in Zhangpu, Fujian; Figure S3: Streak plate isolation of pathogenic bacteria from various tissues of diseased fish in Raoping, Guangdong; Table S1: Bacterial diversity and abundance detected in *Neobenedenia girellae* using 16s rRNA amplicon sequencing; Table S2: Bacterial diversity and abundance detected in *Zeylanicobdella arugamensis* using 16s rRNA amplicon sequencing; Table S3: Bacterial diversity and abundance detected in *Limnotrachelobdella okae* using 16s rRNA amplicon sequencing; Supplementary file 1: Sequences of pathogens isolated from diseased fish in Ningde; Supplementary file 2: Sequences of pathogens isolated from diseased fish in Zhangpu; Supplementary file 3: Sequences of pathogens isolated from diseased fish in Raoping; Video S1: Low-magnification microscopic observation of *N. girellae* ingesting blood containing *T. carassii spectrum*; Video S2: Motile *T. carassii spectrum* observed within *N. girellae* one hour post-feeding.

## Author Contributions

Conceptualization, J.C., S.H. and D.H.L.; methodology, J.C., S.H. and D.H.L.; validation, J.C., J.Z., X.L. and M.L.; formal analysis, J.C. and J.Z.; investigation, J.C., J.Z., Q.L. and N.Y.; resources, J.C., D.H.L. and S.H.; data curation, J.Z. and J.C.; writing-original draft preparation, J.C.; writing-review and editing, S.H. and D.H.L.; visualization, J.Z. and J.C.; supervision, S.H.; project administration, J.C.; funding acquisition, J.C. and S.H.. All authors have read and agreed to the published version of the manuscript.

## Acknowledgments

This work was financially supported by the National Key Research and Development Program (2025YFD2400300), Guangdong S&T Program (2026B0202170002) and National Natural Science Foundation of China (32373171) to Jinsong Chen in Sun Yat-sen University, Guangzhou, China.

## Ethics statement

All animal procedures were approved by the Laboratory Animal Use and Care Committee of Sun Yat-Sen University (license number 32170470).

