## Supplementary Materials and Table 2 for "Parasitic Vectors in Aquaculture: *Neobenedenia girellae* and Leeches as Potential Transmission Agents for *Trypanosoma carassii spectrum* and Pathogenic bacteria": Supplementary file 1_Sequences of the Pathogens Isolated from Ningde.docx

>*Neobenedenia girellae*

TCAGTAAGCGGTGGAAAGCAAACTAAACAGGATTCTCTTAGTAACTGCGAGTGAACAGAGATTAGCCCAACATCGAAGCCACCATGCATTTGTATGGGAGGCAATGTGATGTTTTGGTGTTTGAACGCACTGTCTGGCTAACTTGAAGTCCATATTTGACTGTGGCTTGTATTTATACACCCAGAGATGGTGATAGGCCCGTTCAGGTTGGTATTCTGGCATGACAGTTCAGTGTGTAATCATTCATCATAGAGTCGGATTGCTTGAGAGTGCAGTCCAAAATGGGTGGTAAACTCCATCTAAGGCTAAATACCTGTACGAGTCCGATAGTCAACAAGTACCGTGAGGGAAAGTTGAAAAGTACTCTGAAGAGAGAGTAAATAGTACGTGAAACCACATGGAGGTTAAACAAATAGAGTCAAGCTGCTTGCATCATATCATTGCAATATGCGCATTTTGTTGCATGACATTTGCCGGGACCTGGCGTGTGATTCTGTCTACTTGTAGGCATTTGAACATGTGTTTGTTGTTTGAAAGGTCTCGGTGAGTGTCTGCGCTTATTGTTCATGATGGGTGCTAATGCCATAGTCGGTTGTACTACCAGTGCGCAGTAATCGGCGATTGTGCGGGCTGACTTATTGTCGTTCGTGCAATCAGTGAAACAGTTGGCAACAGTGTGTGCTTGCATTCGCTTTTGTCAAAGAAGGCCACGGCCTGTTGATTACTGTGAACATTGTGAGAGTCGACCGTTCATATGTGCTCATATAATTTGTGGTAGTAAGTCGACTTGTCGACTACTTCCATAAATGAGCTTATGAGCTATGGTATTGTGGTAGACTTTCTATTTGACC

> *Trypanosoma carassii spectrum*

CTGCACTTGCAGGATCTGCGCATGGCTCATTACATCAGACGTAATCTGCCGCAAACATGTTGCGGTTTCCGCATTATTGGATAACTTGGCGAAACGCCAAGCTAATACATGAGTAGAAGGAACGTTCTCTGTTTCGGGTGGTGGGGCAACTCACTGCTTATGGGACGTTCAGCGAATGAATGAAAGTAAAACCAATGCCCTTTTTGGGCAGTATCACCTAGAAGTGTTGACTCAATTCATTCCGTGCGAAAGCCGGATTTTCCGGCGTCTTTTGACGAACAACTGCCCTATCAGCCAGTGATGGCCGTGTAGTGGACTGCCATGGCGTTGACGGGAGCGGGGGATTAGGGTTCGATTCCGGAGAGGGAGCCTGAGAAATAGCTACCACTTCTACGGAGGGCAGCAGGCGCGCAAATTGCCCAATGTCAAAACAAACGATGAGGCAGCGAAAAGAAATAGAGCCGACAGTCCCATCTGGGATTGTCGTATTCAATGGGGGTTATTTAATACCATCCAATATCGAGTAACAATTGGAGGACAAGTCTGGTGCCAGCACCCGCGGTAATTCCAGCTCCAAAAGCGTATATTAATGCTGTTGCTGTTAAAGGGTTCGTAGTTGAATTGAGGGCCTCTGAGGCGCACTGGTATGTCCCGTTCACTTCGAATTTGGTGACCCAGGCCCTTGTGGTCCGTGAACACATTCAGAAACAAGAAACACGGGAGTGGTTCCCTTCCTGATTTTCGCATGTCATGCATGCCAGGGGGCGCCCGTGATTTTTTACTGTGACTAAAAAAGTGTGACCAAAGCAGTCATCCGACTTGAATTAGAAAGCATGGGATAACAAAGGAGCAGCCTATGAGCTACCGTTTCGGCTTTTGTTGGTTTTAAAAACTCATTGGAGATTATGGAGTTGTGCGACAAGCGGCCGGGCGCCTAGTTTTATCTGGTGCCCGTCGCCTTTGTGGGAAACCCCGTACCGATGCAGGAGGGAGGCGGGCTTCGGCTTGTCTTTTCCTCTCGGTGCATTCCCTCAACTCACTGCTTCCAGGAATGAAGGAGGGTAGTTCGGGGGAGAACGTACTGGTGCGTCAGAGGTGAAATTCTTAGACCGCACCAAGACGAACTACAGCGAAGGCATTCTTCAAGGATACCTTCCTCAATCAAGAACCAAAGTGTGGGGATCGAAGATGATTAGAGACCATTGTAGTCCACACTGCAAACGATGACACCCATGAATTGGGGAGTTTTTGGTCGCAGGCGGTGTCGGGTTCATCTCGCTCATCGTCTCACCAATGATATCAATTTACGTGCATATTCTTTTTTCGGTCCCCGCAAGGGGGCTTTTTACGGGAATATCCTCAGCACGTTTTCTTACTTCTTCACGCGAAAGCTTTGAGGTTACAGTCTCAGGGGGGAGTACGTTCGCAAGAGTGAAACTTAAAGAAATTGACGGAATGGCACCACAAGACGTGGAGCGTGCGGTTTAATTTGACTCAACACGGGGAACTTTACCAGATCCGGACAGGGTGAGGATTGACAGATTGAGTGTTCTTTCTCGATCCCCTGAATGGTGGTGCATGGCCGCTTTTGGTCGGTGGAGTGATTTGTTTGGTTGATTCCGTCAACGGACGAGATCCAAGCTGCCCAGTAGGATTCAGAATTGCCCATAGGATAGCAATCCCTTCCGCGGGTTTTACCCTAAGGGGGGGCGGTATTCGTTTGTATCCTTCTCTGCGGGATTCCTTGACTTTGCACAAGGTGAGATTTTGGGCAACAGCAGGTCTGTGATGCTCCTCAATGTTCTGGGCGACACGCGCACTACAATGTCAGTGAGAACAAGAAAAACGACTCTTGTCGTACCTACTTGATCAAAAGAGTGGGAAAACACCGGAATCACATAGACCCACTTGGGACCGAGTATTGCAATTATTGGTCGCGCAACGAGGAATGTCTCGTAGGCGCAGCTCATCAAACTGTGCCGATTACGTCCCTGCCATTTGTACACACCGCCCGTCGTTGTTTCCGATGATGGTACAATACAGGTGATCGGACAGTCGAGTGTCTCACTG

> *T. carassii spectrum* from *N. girellae* _1

TCTATGCTTGTTTCAAGGACTTAGCCATGCATGCCTCAGAATCACTGCACTTGCAGGAATCTGCGCATGGCTCATTACATCAGACGTAATCTGCCGCAAACATGTTGCGGTTTCCGCATTATTGGATAACTTGGCGAAACGCCAAGCTAATACATGAGTAGAAGGAACGTTCTCTGTTTCGGGTGGTGGGGCAACTCACTGCTTATGGGACGTTCAGCGAATGAATGAAAGTAAAACCAATGCCCTTTTTGGGCAGTATCACCTAGAAGTGTTGACTCAATTCATTCCGTGCGAAAGCCGGATTTTCCGGCGTCTTTTGACGAACAACTGCCCTATCAGCCAGTGATGGCCGTGTAGTGGACTGCCACGGCGTTGACGGGAGCGGGGGATTAGGGTTCGATTCCGGAGAGGGAGCCTGAGAAATAGCTACCACTTCTACGGAGGGCAGCAGGCGCGCAAATTGCCCAATGTCAAAACAAACGATGAGGCAGCGAAAAGAAATAGAGCCGACAGTCCCATCTGGGATTGTCGTATTCAATGGGGGTTATTTAATACCATCCAATATTGAGTAACAATTGGAGGACAAGTCTGGTGCCAGCACCCGCGGTAATTCCAGCTCCAAAAGCGTATATTAATGCTGTTGCTGTTAAAGGGTTCGTAGTTGAATTGAGGGCCTCTGAGGCGCACTGGTATGTCCCGTTCACTTCGAATTTGGTGACCCAGGCCCTTGTGGTCCGTGAACACATTCAGAAACAAGAAACACGGGAGTGGTTCCCTTCCCGATTTTCGCATGTCATGCATGCCAGGGAGCGCCCGTGATTTTTTACTGTGACTAAAAAAGTGTGACCAAAGCAGTCATCCGACTTGAATTAGAAAGCATGGGATAACAAAGGAGCAGCCTATGAGCTACCGTTTCGGCTTTTGTTGGTTTTAAAAACTCATTGGAGATTATGGAGTTGTGCGACAAGCGGCCGGGCGCCTAGTTTTATCTGGTGCCCGCCGCCTTTGTGGGAAACCCCGTACCGATGCAGGAGGGAGGCGGGCTTCGGCTTGTCTTTTCCTCTCGGTGCATTCCCTCAACTCACTGCTTCCAGGAATGAAGGAGGGTAGTTCGGGGGAGAACGTACTGGTGCGTCAGAGGTGAAATTCTTAGACCGCACCAAGACGAACTACAGCGAAGGCATTCTTCAAGGATACCTTCCTCAATCAAGAACCAAAGTGTGGGGATCGAAGATGATTAGAGACCATTGTAGTCCACACTGCAAACGATGACACCCATGAATTGGGGAGTTTTTGGTCGCAGGCGGTGTCGGGTTCGTCTCGCTCATCGTCTCACCAATGATATCAATTTACGTGCATATTCTTTTTTCGGTCCCCGCAAGGGGGCTTTTTACGGGAATATCCTCAGCACGTTTTCTTACTTCTTCACGCGAAAGCTTTGAGGTTACAGTCTCAGGGGGGAGTACGTTCGCAAGAGTGAAACTTAAAGAAATTGACGGAATGGCACCACAAGACGTGGAGCGTGCGGTTTAATTTGACTCAACACGGGGAACTTTACCAGATCCGGACAGGGTGAGGATTGACAGATTGAGTGTTCTTTCTCGATCCCCTGAATGGTGGTGCATGGCCGCTTTTGGTCGGTGGAGTGATTTGTTTGGTTGATTCCGTCAACGGACGAGATCCAAGCTGCCCAGTAGGATTCAGAATTGCCCATAGGATAGCAATCCCTTCCGCGGGTTTTACCCTAAGGGGGGGCGGTATTCGTTTGTATCCTTCTCTGCGGGATTCCTTGACTTTGCACAAGGTGAGATTTTGGGCAACAGCAGGTCTGTGATGCTCCTCAATGTTCTGGGCGACACGCGCACTACAATGTCAGTGAGAACAAGAAAAACGACTCTTGTCGTACCTACTTGATCAAAAGAGTGGGAAAACACCGGAATCACATAGACCCACTTGGGACCGAGTATTGCAATTATTGGTCGCGCAACGAGGAATGTCTCGTAGGCGCAGCTCATCAAACTGTGCCGATTACGTCCCTGCCATTTGTACACACCGCCCGTCGTTGTTTCCGATGATGGTACAATACAGGTGATCGGACAGTCGAGTGTCTCACTTGACTGAAAGTTCACCGATATTGTATCAATAGAGGAAGCAAAAGTCAATCTCTAGAGGTCCCCGGGTA

> *T. carassii spectrum* from *N. girellae* _2

TCTATGCTTGTTTCAAGGACTTAGCCATGCATGCCTCAGAATCACTGCACTTGCAGGAATCTGCGCATGGCTCATTACATCAGACGTAATCTGCCGCAAACAAGTTGCGGTTTCCGCATTATTGGATAACTTGGCGAAACGCCAAGCTAATACATGAGTAGAAGGAACGTTCTCTGTTTCGGGTGGTGGGGCAACTCACTGCTTATGGGACGTTCAGCGAATGAATGAAAGTAAAACCAATGCCCTTTTTGGGCAGTATCACCTAGAAGTGTTGACTCAATTCATTCCGTGCGAAAGCCGGATTTTCCGGCGTCTTTTGACGAACAACTACCCTATCAGCCAGTGATGGCCGTGTAGTGGACTGCCATGGCGTTGACGGGAGCGGGGGATTAGGGTTCGATTCCGGAGAGGGAGCCTGAGAAATAGCTACCACTTCTACGGAGGGCAGCAGGCGCGCAAATTGCCCAATGTCAAAACAAACGATGAGGCAGCGAAAAGAAATAGAGCCGACAGTCCCATCTGGGGTTGTCGTATTCAATGGGGGTTATTTAATACCATCCAATATCGAGTAACAATTGGAGGACAAGTCTGGTGCCAGCACCCGCGGTAATTCCGGCTCCAAAAGCGTATATTAATGCTGTTGCTGTTAAAGGGTTCGTAGTTGAATTGAGGGCCTCTGAGGCGCACTGGTATGTCCCGTTCACTTCGAATTTGGTGACCCAGGCCCTTGTGGTCCGTGAACACATTCAGAAACAAGAAACACGGGAGTGGTTCCCTTCCTGATTTTCGCATGTCATGCATGCCAGGGGGCGCCCGTGATTTTTTACTGTGACTAAAAAAGTGTGACCAAAGCAGTCATCCGACTTGAATTAGAAAGCATGGGATAACAAAGGAGCAGCCTATGAGCTACCGTTTCGGCTTTTGTTGGTTTTAAAAACTCGTTGGAGATTATGGAGTTGTGCGACAAGCGGCCGGGCGCCTAGTTTTATCTGGTGCCCGTCGCCTTTGTGGGAAACCCCGTACCGATGCAGGAGGGAGGCGGGCTTCGGCTTGTCTTTTCCTCTCGGTGCATTCCCTCAACTCACTGCTTCCAGGAATGAAGGAGGGTAGTTCGGGGGAGAACGTACTGGTGCGTCAGAGGTGAAATTCTTAGACCGCACCAAGACGAACTACAGCGAAGGCATTCTTCAAGGATACCTTCCTCAATCAAGAACCAAAGTGTGGGGATCGAAGATGATTAGAGACCATTGTAGTCCACACTGCAAACGATGACACCCATGAATTGGGGAGTTTTTGGTCGCAGGCGGTGTCGGGTTCATCTCGCTCATCGTCTCACCAATGATATCAATTACGTGCATATTCTTTTTTCGGTCCCCGCAAGGGGCTTTTTACGGGAATATCCTCAGCACGTTTTCTTACTTCTTCACGCGAAAGCTTTGAGGTTACAGTCTCAGGGGGGGGGTACGTTCGCAAGAGTAAAACTTAAAGAAATTGACGGAATGGCACCACAAGACGTGGAGCGTGCGGTTTAATTTGACTCAACACGGGGAACTTTACCAGATCCGGACAGGGTGAGGATTGACAGATTGAGTGTTCTTTCTCGATCCCCTGAATGGTGGTGCATGGCCGCTTTTGGTCGGTGGAGTGATTTGTTTGGTTGATTCCGTCAACGGACGAGATCCAAGCTGCCCAGTAGGATTCAGAATTGCCCATAGGATAGCAATCCCTTCCGCGGGTTTTACCCTAAGGGGGGGCGGTATTCGTTTGTATCCTTCTCTGCGGGATTCCTTGACTTTGCACAAGGTGAGATTTTGGGCAACAGCAGGTCTGTGATGCTCCTCAATGTTCTGGGCGACACGCGCACTACAATGTCAGTGAGAACAAGAAAAACGACTCTTGTCGTACCTACTTGATCAAAAGAGTGGGAAAACACCGGAATCACATAGACCCACTTGGGACCGAGTATTGCAATTATTGGTCGCGCAACGAGGAATGTCTCGTAGGCGCAGCTCATCAAACTGTGCCGATTACGTCCCTGCCATTTGTACACACCGCCCGTCGTTGTTTCCGATGATGGTACAATACAGGTGATCGGACAGTCGAGTGTCTCACTTGACTGAAAGTTCACCGATATTGTTTCAATAGAGGAAGCAAAAGTC

>*Bacillus cereus*

TTAGGCGGCTGGCTCCAAAAGGTTACCCCACCGACTTCGGGTGTTACAAACTCTCGTGGTGTGACGGGCGGTGTGTACAAGGCCCGGGAACGTATTCACCGCGGCATGCTGATCCGCGATTACTAGCGATTCCAGCTTCATGTAGGCGAGTTGCAGCCTACAATCCGAACTGAGAACGGTTTTATGAGATTAGCTCCACCTCGCGGTCTTGCAGCTCTTTGTACCGTCCATTGTAGCACGTGTGTAGCCCAGGTCATAAGGGGCATGATGATTTGACGTCATCCCCACCTTCCTCCGGTTTGTCACCGGCAGTCACCTTAGAGTGCCCAACTTAATGATGGCAACTAAGATCAAGGGTTGCGCTCGTTGCGGGACTTAACCCAACATCTCACGACACGAGCTGACGACAACCATGCACCACCTGTCACTCTGCTCCCGAAGGAGAAGCCCTATCTCTAGGGTTTTCAGAGGATGTCAAGACCTGGTAAGGTTCTTCGCGTTGCTTCGAATTAAACCACATGCTCCACCGCTTGTGCGGGCCCCCGTCAATTCCTTTGAGTTTCAGCCTTGCGGCCGTACTCCCCAGGCGGAGTGCTTAATGCGTTAACTTCAGCACTAAAGGGCGGAAACCCTCTAACACTTAGCACTCATCGTTTACGGCGTGGACTACCAGGGTATCTAATCCTGTTTGCTCCCCACGCTTTCGCGCCTCAGTGTCAGTTACAGACCAGAAAGTCGCCTTCGCCACTGGTGTTCCTCCATATCTCTACGCATTTCACCGCTACACATGGAATTCCACTTTCCTCTTCTGCACTCAAGTCTCCCAGTTTCCAATGACCCTCCACGGTTGAGCCGTGGGCTTTCACATCAGACTTAAGAAACCACCTGCGCGCGCTTTACGCCCAATAATTCCGGATAACGCTTGCCACCTACGTATTACCGCGGCTGCTGGCACGTAGTTAGCCGTGGCTTTCTGGTTAGGTACCGTCAAGGTGCCAGCTTATTCAACTAGCACTTGTTCTTCCCTAACAACAGAGTTTTACGACCCGAAAGCCTTCATCACTCACGCGGCGTTGCTCCGTCAGACTTTCGTCCATTGCGGAAGATTCCCTACTGCTGCCTCCCGTAGGAGTCTGGGCCGTGTCTCAGTCCCAGTGTGGCCGATCACCCTCTCAGGTCGGCTACGCATCGTTGCCTTGGTGAGCCGTTACCTCACCAACTAGCTAATGCGACGCGGGTCCATCCATAAGTGACAGCCGAAGCCGCCTTTCAATTTCGAACCATGCGGTTCAAAATGTTATCCGGTATTAGCCCCGGTTTCCCGGAGTTATCCCAGTCTTATGGGCAGGTTACCCACGTGTTACTCACCCGTCCGCCGCTAACTTCATAAGAGCAAGCTCTTAATCCATTCGCTCGACTTGCAGTATAGG

>*Photobacterium damselae*

CCCTCCCGAAGGTTAAGCTATCTACTTCTGGTGCAGCCCACTCCCATGGTGTGACGGGCGGTGTGTACAAGGCCCGGGAACGTATTCACCGTGGCATTCTGATCCACGATTACTAGCGATTCCGACTTCACGGAGTCGAGTTGCAGACTCCGATCCGGACTACGACATACTTTCTGGGATTCGCTCCACCTCGCGGTCTTGCTGCCCTCTGTATATGCCATTGTAGCACGTGTGTAGCCCTACTCGTAAGGGCCATGATGACTTGACGTCGTCCCCACCTTCCTCCGGTTTATCACCGGCAGTCTCCCTGGAGTTCCCACCCGAAGTGCTGGCAAACAAGGATAAGGGTTGCGCTCGTTGCGGGACTTAACCCAACATTTCACAACACGAGCTGACGACAGCCATGCAGCACCTGTCTCAGAGTTCCCGAAGGCACACTCGAATCTCTTCAAGCTTCTCTGGATGTCAAGAGTAGGTAAGGTTCTTCGCGTTGCATCGAATTAAACCACATGCTCCACCGCTTGTGCGGGCCCCCGTCAATTCATTTGAGTTTTAATCTTGCGACCGTACTCCCCAGGCGGTCTACTTAACGCGTTAGCTCCGAAAGCCACGGCTCAAGGCCACAACCTCCAAGTAGACATCGTTTACGGCGTGGACTACCAGGGTATCTAATCCTGTTTGCTCCCCACGCTTTCGCATCTGAGCGTCAGTCTTTGTCCAGGGGCCGCCTTCGCCACTGGTATTCCTTCAGATCTCTACGCATTTCACCGCTACACCTGAAATTCTACCCCCCTCTACAAGACTCTAGCCTGCCAGTTTCAAATGCAATTCCGAGGTTAAGCCCCGGGCTTTCACATCTGACTTAACAGGCCGCCTGCATGCGCTTTACGCCCAGTAATTCCGATTAACGCTCGCACCCTCCGTATTACCGCGGCTGCTGGCACGGAGTTAGCCGGTGCTTCTTCTGTAGGTAACGTCAAATAATGCAGGTGTTAACTACACTACCTTCCTCCCTACTGAAAGTGCTTTACAACCCGAAGGCCTTCTTCACACACGCGGCATGGCTGCATCAGGGTTTCCCCCATTGTGCAATATTCCCCACTGCTGCCTCCCGTAGGAGTCTGGACCGTGTCTCAGTTCCAGTGTGGCTGATCATCCTCTCAGACCAGCTAGGGATCGTTGCCTTGGTGAGCCATTACCTCACCAACAAGCTAATCCCACCTGGGCTAATCCTGACGCGAGAGGCCCGAAGGTCCCCCTCTTTGCTCCGAAGAGATTATGCGGTATTAGCTATCGTTTCCAATAGTTATCCCCCACATCAGGGCATATTCCCAGGCATTACTCACCCGTCCGCCGCTCGCCGCCCTTAACGTTCCCCGAAGGTTCAGTTAAGTCGCTGCCGCTCGACT

>*Bacillus thuringiensis*

ACGACTTACTGAGCCAGGATCAAACTCTATGGTTACCTTGTTACGACTTACTGAGCCAGGATCAAACTCTATGGTTACCTTGTTACGACTTACTGAGCCAGGATCAAACTCTATGGTTACCTTGTTACGACTTACTGAGGCAGGATCAAACTCTATGGGTACCTTGTTACGACTTATTGAGTCCGGCTCAAAGTCTAGGGGTTCTTTGTACCGACCATTGGAGCACGTGTGTAACCCAAGGCATAAGGGGCATGATGATTTGACGTCATCCCCACCTTCCTCCGGTTTGTCACCGGCAGTCACCTTAGAGTGCCCAACTTAATGATGGCAACTAAGATCAAGGGTTGCGCTCGTTGCGGGACTTAACCCAACATCTCACGACACGAGCTGACGACAACCATGCACCACCTGTCACTCTGCTCCCGAAGGAGAAGCCCTATCTCTAGGGTTTTCAGAGGATGTCAAGACCTGGTAAGGTTCTTCGCGTTGCTTCGAATTAAACCACATGCTCCACCGCTTGTGCGGGCCCCCGTCAATTCCTTTGAGTTTCAGCCTTGCGGCCGTACTCCCCAGGCGGAGTGCTTAATGCGTTAACTTCAGCACTAAAGGGCGGAAACCCTCTAACACTTAGCACTCATCGTTTACGGCGTGGACTACCAGGGTATCTAATCCTGTTTGCTCCCCACGCTTTCGCGCCTCAGTGTCAGTTACAGACCAGAAAGTCGCCTTCGCCACTGGTGTTCCTCCATATCTCTACGCATTTCACCGCTACACATGGAATTCCACTTTCCTCTTCTGCACTCAAGTCTCCCAGTTTCCAATGACCCTCCACGGTTGAGCCGTGGGCTTTCACATCAGACTTAAGAAACCACCTGCGCGCGCTTTACGCCCAATAATTCCGGATAACGCTTGCCACCTACGTATTACCGCGGCTGCTGGCACGTAGTTAGCCGTGGCTTTCTGGTTAGGTACCGTCAAGGTGCCAGCTTATTCAACTAGCACTTGTTCTTCCCTAACAACAGAGTTTTACGACCCGAAAGCCTTCATCACTCACGCGGCGTTGCTCCGTCAGACTTTCGTCCATTGCGGAAGATTCCCTACTGCTGCCTCCCGTAGGAGTCTGGGCCGTGTCTCAGTCCCAGTGTGGCCGATCACCCTCTCAGGTCGGCTACGCATCGTTGCCTTGGTGAGCCGTTACCTCACCAACTAGCTAATGCGACGCGGGTCCATCCATAAGTGACAACAGTATCTGCCTACCAATTTACGACCACTGAGCCAGGATCAAACTCTATGGTTACCTTGTTACGACTTACTGAGCCAGGATCAAACTCTATGGTTACCTTGTTACGACTTACTGAGCCAGGATCAAACTCTATGGTTACCTTGTTACGACTTACTGAGCCAGGATCAAACTC

>*Vibrio harveyi*

TACCTTGTTACGACTTACTGAGCCAGGATCAAACTCTATGGTTACCTTGTTACGACTTACTGAGCCAGGATCAAACGCTATGGTTACCTTGTTACGAGTTACTGAGCCAGGATCAAACTCTATGGTTACCTTTCTCCGAGTTAAGTTGCAAAGTCCGATCCGGACTACGACGCACTTTTTGGGATTCGCTCACTCTCGCAAGTTGGCCGCCCTCTGTATGCGCCATTGTAGCACGTGTGTAGCCCTACTCGTAAGGGCCATGATGACTTGACGTCGTCCCCACCTTCCTCCGGTTTATCACCGGCAGTCTCCCTGGAGTTCCCACCCGAAGTGCTGGCAAACAAGGATAAGGGTTGCGCTCGTTGCGGGACTTAACCCAACATTTCACAACACGAGCTGACGACAGCCATGCAGCACCTGTCTCAGAGTTCCCGAAGGCACCAATCCATCTCTGGAAAGTTCTCTGGATGTCAAGAGTAGGTAAGGTTCTTCGCGTTGCATCGAATTAAACCACATGCTCCACCGCTTGTGCGGGCCCCCGTCAATTCATTTGAGTTTTAATCTTGCGACCGTACTCCCCAGGCGGTCTACTTAACGCGTTAGCTCCGAAAGCCACGGCTCAAGGCCACAACCTCCAAGTAGACATCGTTTACGGCGTGGACTACCAGGGTATCTAATCCTGTTTGCTCCCCACGCTTTCGCATCTGAGTGTCAGTATCTGTCCAGGGGGCCGCCTTCGCCACCGGTATTCCTTCAGATCTCTACGCATTTCACCGCTACACCTGAAATTCTACCCCCCTCTACAGTACTCTAGTCTGCCAGTTTCAAATGCTATTCCGAGGTTGAGCCCCGGGCTTTCACATCTGACTTAACAAACCACCTGCATGCGCTTTACGCCCAGTAATTCCGATTAACGCTCGCACCCTCCGTATTACCGCGGCTGCTGGCACGGAGTTAGCCGGTGCTTCTTCTGTCGCTAACGTCAAATAATGCAGCTATTAACTACACTACCTTCCTCACGACTGAAAGTGCTTTACAACCCGAAGGCCTTCTTCACACACGCGGCATGGCTGCATCAGGCTTGCGCCCATTGTGCAATATTCCCCACTGCTGCCTCCCGTAGGAGTCTGGACCGTGTCTCAGTTCCAGTGTGGCTGATCATCCTCTCAGACCAGCTAGGGATCGTCGCCTTGGTGAGCCATTACCTCACCAACTAGCTAATCCCACCTAGGCATATCCTGACGCGAGAGGCCCGAAGGTCCCCCTCTTTGACCCGTAGGTATTATGCGGTATTAGCCATCGTTTCCAATGGTTATCCCCCACACCAGGGCAATTACACTGTATGGTTTCCTTGTTACGACTTACTGAGCCAGGATCAAACTCTATGGTTACCTTGTTACGACTTACTGAGCCAGGATCAAACTCTA

>*Priestia megaterium*

TACCTTGTTACGACTTACTGAGCCAGGATCAAACTCTATGGTTACCTTGTTACGACTTACTGAGCCAGGATCAAACTCTATGGTTACCTTGTTACGACTTACTGAGCCAGGATCAAACTCTATGGTTACCTTGTTACGACTTACTGAGCCAGGATCAAACTCTATGGTTACCTTGTTACGACTTACTGAGCCAGGATCAAACTCTATGGTTACCTTGTTACGACTTACTGAGCCAGGATCAAACTCTATGGTTACCTTGTTACGACTTACTGAGCCTGGACCCCTTCCTATGGTTTGCCTCCGGCGACCACCTTAGAGTGGCCAACTAAATGCTGGCAACTAAGATCAAGGGTTGCGCTCGTTGCGGGACTTAACCCAACATCTCACGACACGAGCTGACGACAACCATGCACCACCTGTCACTCTGTCCCCCGAAGGGGAACGCTCTATCTCTAGAGTTGTCAGAGGATGTCAAGACCTGGTAAGGTTCTTCGCGTTGCTTCGAATTAAACCACATGCTCCACCGCTTGTGCGGGCCCCCGTCAATTCCTTTGAGTTTCAGTCTTGCGACCGTACTCCCCAGGCGGAGTGCTTAATGCGTTAGCTGCAGCACTAAAGGGCGGAAACCCTCTAACACTTAGCACTCATCGTTTACGGCGTGGACTACCAGGGTATCTAATCCTGTTTGCTCCCCACGCTTTCGCGCCTCAGCGTCAGTTACAGACCAAAAAGCCGCCTTCGCCACTGGTGTTCCTCCACATCTCTACGCATTTCACCGCTACACGTGGAATTCCGCTTTTCTCTTCTGCACTCAAGTTCCCCAGTTTCCAATGACCCTCCACGGTTGAGCCGTGGGCTTTCACATCAGACTTAAGAAACCGCCTGCGCGCGCTTTACGCCCAATAATTCCGGATAACGCTTGCCACCTACGTATTACCGCGGCTGCTGGCACGTAGTTAGCCGTGGCTTTCTGGTTAGGTACCGTCAAGGTACGAGCAGTTACTCTCGTACTTGTTCTTCCCTAACAACAGAGTTTTACGACCCGAAAGCCTTCATCACTCACGCGGCGTTGCTCCGTCAGACTTTCGTCCATTGCGGAAGATTCCCTACTGCTGCCTCCCGTAGGAGTCTGGGCCGTGTCTCAGTCCCAGTGTGGCCGATCACCCTCTCAGGTCGGCTATGCATCGTTGCCTTGGTGAGCCGTTACCTCACCTATAAGGTAACCCTCCGCGGGCCCATCTGTAAAGGATAAAACTCTATGGTTACCTTGTTACGACTTACTGAGCCAGGATCAAACTCTATGGTTACCTTGTTACGACTTACTGAGCCAGGATCAAACTCTATGGTTACCTTGTTACGACTTACTGAGCCAGGATC

>*Psychrobacter celer*

CCTCACTAAGTTAGGCTAACCACTTCTGGTGCAATCAACTCCCATGGTGTGACGGGCGGTGTGTACAAGGCCCGGGAACGTATTCACCGCGGCATTCTGATCCGCGATTACTAGCGATTCCTACTTCATGGAGTCGAGTTGCAGACTCCAATCTGGACTACGATAGGCTTTTTGAGATTCGCATCACATCGCTGTGTAGCTGCCCTCTGTACCTACCATTGTAGCACGTGTGTAGCCCTGGTCGTAAGGGCCATGATGACTTGACGTCGTCCCCGCCTTCCTCCAGTTTGTCACTGGCAGTATCCTTAGAGTTCCCGGCTTAACCCGCTGGTAACTAAGGACAAGGGTTGCGCTCGTTGCGGGACTTAACCCAACATCTCACGACACGAGCTGACGACAGCCATGCAGCACCTGTATCACAATTCCCGAAGGCACTCTCGTATCTCTACAAGATTCTGTGTATGTCAAGACCAGGTAAGGTTCTTCGCGTTGCATCGAATTAAACCACATGCTCCACCGCTTGTGCGGGCCCCCGTCAATTCATTTGAGTTTTAACCTTGCGGCCGTACTCCCCAGGCGGTCTACTTATTGCGTTAGCTGCGTCACTAAGTCCTCAAGGGACCCAACGACTAGTAGACATCGTTTACGGCGTGGACTACCAGGGTATCTAATCCTGTTTGCTACCCACGCTTTCGAACCTCAGTGTCAGTATGATGCCAGGAAGCTGCCTTCGCCATCGGTATTCCTCCAGATCTCTACGCATTTCACCGCTACACCTGGAATTCTACTTCCCTCTCACCTACTCTAGCTTAACAGTATCAGATGCAGTTCCCAGGTTAAGCCCGGGGATTTCACATCTGACTTATCAAGCCACCTACGCTCGCTTTACGCCCAGTAATTCCGATTAACGCTTGCACCCTCTGTATTACCGCGGCTGCTGGCACAGAGTTAGCCGGTGCTTATTCTGCAGCTAATGTCATCGCTAACGGGTATTAACCGTTAGGTCTTCTTCACTGCTTAAAGTGCTTTACAACCAAAAGGCCTTCTTCACACACGCGGCATGGCTGGATCAGGGTTTCCCCCATTGTCCAATATTCCCCACTGCTGCCTCCCGTAGGAGTCCGGGCCGTGTCTCAGTCCCGGTGTGGCTGATCATCCTCTCAGACCAGCTACAGATCGTCGCCATGGTAGGCCTTTACCCCACCATCTAGCTAATCCGACTTAGGCTCATCTAATAGCGAGAGCAGTAAACTGCCCCCTTTCTCCCGTAGGACGTATGCGGTATTAATTCGAGTTTCCCCGAGCTATCCCCCACTACTAGGTAGATTCCTAAGTATTACTCACCCGTCCGCCGCTCGACGCCTGGGAGCAAGCTCCCATCGTTTCCGCTCGACTTGCAGGTAAGCTT
