## Supplementary Materials and Table 2 for "Parasitic Vectors in Aquaculture: *Neobenedenia girellae* and Leeches as Potential Transmission Agents for *Trypanosoma carassii spectrum* and Pathogenic bacteria": Supplementary file 2_Sequences of the Pathogens Isolated from Zhangpu.docx

> *Zeylanicobdella arugamensis*

AAGATATTGGAACTTTATATTTTATATTCGGAGCTTGATCTGCAATAATTGGAACTAGTATAAGGTTTGTAATTCGAGCAGAATTATCACAACCAGGTAGATTTATAGAAAATGATCAAATTTATAACTCAATAATTACAGCACATGGGTTAATTATAATTTTTTTTATAGTTATACCAATTCTAATTGGTGGTTTTGGTAACTGGCTAGTACCTTTAATAATTGGTGCTCCAGACATGGCTTTTCCACGACTTAATAATCTTAGATTTTGATTATTACCACCATCATTAATTCTTCTTTTATCATCTGCATTAATTGAAAGTGGGGTTGGTACAGGTTGAACAGTCTACCCACCATTATCAGCTAATATAGCTCATTCTGGACCATCAGTAGATTTAGCAATTTTTTCTTTACATTTAGCAGGAATTTCATCAATTTTAGGTGCATTAAATTTTATTACAACAATTTTAAATATACGATGAAATGGGTTAAAATTAGAACGTATATCATTATTTGTTTGAGCAGTATTAATTACTGTTGTATTATTACTTTTATCTTTACCTGTATTAGCAGCTGCTATTACAATATTACTTACAGATCGTAATTTTAATACATCATTCTTCGATCCAGTTGGTGGTGGAGATCCAATTTTATACCAACATTTATTTTGATTTTTTGG

>*Vibrio campbellii*

GTGGGGAATATTGCACAATGGGCGCAAGCCTGATGCAGCCATGCCGCGTGTGTGAAGAAGGCCTTCGGGTTGTAAAGCACTTTCAGTCGTGAGGAAGGTAGTGTAGTTAATAGCTGCATTATTTGACGTTAGCGACAGAAGAAGCACCGGCTAACTCCGTGCCAGCAGCCGCGGTAATACGGAGGGTGCGAGCGTTAATCGGAATTACTGGGCGTAAAGCGCATGCAGGTGGTTTGTTAAGTCAGATGTGAAAGCCCGGGGCTCAACCTCGGAACTGCATTTGAAACTGGCAAACTAGAGTACTGTAGAGGGGGGTAGAATTTCAGGTGTAGCGGTGAAATGCGTAGAGATCTGAAGGAATACCAGTGGCGAAGGCGGCCCCCTGGACAGATACTGACACTCAGATGCGAAAGCGTGGGGAGCGAACAGG

>*Photobacterium damselae*

AGTAACTGGGCGGCGGCTACCTGCAAGTCGAGCGGCAGCGACTTAACTGAACCTTCGGGGAACGTTAAGGGCGGCGAGCGGCGGACGGGTGAGTAATGCCTGGGAATATGCCCTGATGTGGGGGATAACTATTGGAAACGATAGCTAATACCGCATAATCTCTTCGGAGCAAAGAGGGGGACCTTCGGGCCTCTCGCGTCAGGATTAGCCCAGGTGGGATTAGCTTGTTGGTGAGGTAATGGCTCACCAAGGCAACGATCCCTAGCTGGTCTGAGAGGATGATCAGCCACACTGGAACTGAGACACGGTCCAGACTCCTACGGGAGGCAGCAGTGGGGAATATTGCACAATGGGGGAAACCCTGATGCAGCCATGCCGCGTGTGTGAAGAAGGCCTTCGGGTTGTAAAGCACTTTCAGTAGGGAGGAAGGTAGTGTAGTTAACACCTGCATTATTTGACGTTACCTACAGAAGAAGCACCGGCTAACTCCGTGCCAGCAGCCGCGGTAATACGGAGGGTGCGAGCGTTAATCGGAATTACTGGGCGTAAAGCGCATGCAGGCGGCCTGTTAAGTCAGATGTGAAAGCCCGGGGCTTAACCTCGGAATTGCATTTGAAACTGGCAGGCTAGAGTCTTGTAGAGGGGGGTAGAATTTCAGGTGTAGCGGTGAAATGCGTAGAGATCTGAAGGAATACCAGTGGCGAAGGCGGCCCCCTGGACAAAGACTGACGCTCAGATGCGAAAGCGTGGGGAGCAAACAGGATTAGATACCCTGGTAGTCCACGCCGTAAACGATGTCTACTTGGAGGTTGTGGCCTTGAGCCGTGGCTTTCGGAGCTAACGCGTTAAGTAGACCGCCTGGGGAGTACGGTCGCAAGATTAAAACTCAAATGAATTTGACGGGGGCCCGCACAAGCGGTGGAGCATGTGGTTTAATTCGATGCAACGCGAGATCTTACTACTCTGACATCCAGAGAAGCTCGAGAGATTCGAGTGTGCCTTCGGGAACTCTGAGAACAGGTGCTGCATGGCTGTCGTCAGCTC

>*Vibrio ponticus*

TTACCTGGGGGCGGCCTATACTGCAAGTCGAGCGGCAGCGACAACATTGAACCTTCGGGGGATTTGTTGGGCGGCGAGCGGCGGACGGGTGAGTAATGCCTAGGAAATTGCCCTGATGTGGGGGATAACCATTGGAAACGATGGCTAATACCGCATGATGCCTACGGGCCAAAGAGGGGGACCTTCGGGCCTCTCGCGTCAGGATATGCCTAGGTGGGATTAGCTAGTTGGTGAGGTAAGGGCTCACCAAGGCGACGATCCCTAGCTGGTCTGAGAGGATGATCAGCCACACTGGAACTGAGACACGGTCCAGACTCCTACGGGAGGCAGCAGTGGGGAATATTGCACAATGGGCGCAAGCCTGATGCAGCCATGCCGCGTGTATGAAAAAGGCCTTCGGGTTGTAAAGTACTTTCAGTAGGGAGGAAGGTTCATGCGTTAATAGCGTATGGATTTGACGTTACCTACAGAAGAAGCACCGGCTAACTCCGTGCCAGCAGCCGCGGTAATACGGAGGGTGCGAGCGTTAATCGGAATTACTGGGCGTAAAGCGCATGCAGGTGGTTAGTTAAGTCAGATGTGAAAGCCCGGGGCTCAACCTCGGAATTGCATTTGAAACTGGCTGACTAGAGTACTGTAGAGGGGGGTAGAATTTCAGGTGTAGCGGTGAAATGCGTAGAGATCTGAAGGAATACCGGTGGCGAAGGCGGCCCCCTGGACAGATACTGACACTCAGATGCGAAAGCGTGGGGAGCAAACAGGATTAGATACCCTGGTAGTCCACGCCGTAAACGATGTCTACTTGGAGTTGTGGTCTTTGAGCCGTGGCTTTCGGAGCTAACGCGTTAAGTAGACCGCCTGGGGAGTACGGTCGCAAGATTAAAACTCAAATGAATTTGACGGGGGCCCGCACAAGCGGTGGAGCATGTGGTTTAATTCGATGCACGCGAAGATCTTACCTACTCTTGACATCCATAGACTCAGCAGAGATGCTTCTGTGCCTTCGGGACCTATGAGAACAGGTGCTGCATGCTGTCGTCAG

>*Bacillus pumilus*

AGGACTGGCGGCGTGCTATACTGCAAGTCGAGCGGACAGAAGGGAGCTTGCTCCCGGATGTTAGCGGCGGACGGGTGAGTAACACGTGGGTAACCTGCCTGTAAGACTGGGATAACTCCGGGAAACCGGAGCTAATACCGGATAGTTCCTTGAACCGCATGGTTCAAGGATGAAAGACGGTTTCGGCTGTCACTTACAGATGGACCCGCGGCGCATTAGCTAGTTGGTGAGGTAACGGCTCACCAAGGCGACGATGCGTAGCCGACCTGAGAGGGTGATCGGCCACACTGGGACTGAGACACGGCCCAGACTCCTACGGGAGGCAGCAGTAGGGAATCTTCCGCAATGGACGAAAGTCTGACGGAGCAACGCCGCGTGAGTGATGAAGGTTTTCGGATCGTAAAGCTCTGTTGTTAGGGAAGAACAAGTGCAAGAGTAACTGCTTGCACCTTGACGGTACCTAACCAGAAAGCCACGGCTAACTACGTGCCAGCAGCCGCGGTAATACGTAGGTGGCAAGCGTTGTCCGGAATTATTGGGCGTAAAGGGCTCGCAGGCGGTTTCTTAAGTCTGATGTGAAAGCCCCCGGCTCAACCGGGGAGGGTCATTGGAAACTGGGAAACTTGAGTGCAGAAGAGGAGAGTGGAATTCCACGTGTAGCGGTGAAATGCGTAGAGATGTGGAGGAACACCAGTGGCGAAGGCGACTCTCTGGTCTGTAACTGACGCTGAGGAGCGAAAGCGTGGGGAGCGAACAGGATTAGATACCCTGGTAGTCCACGCCGTAAACGATGAGTGCTAAGTGTTAGGGGGTTCCCGCCCCTTAGTGCTGCAGCTAACGCATTAAGCACTCCGCCTGGGGGAGTACGGTCGCAGACTGAAACTCAAAGGA

>*Vibrio harveyi*

ATACCTGGCGGCAGTCTACCTGCAAGTCGAGCGGAAACGAGTTATCTGAACCTTCGGGGAACGATAACGGCGTCGAGCGGCGGACGGGTGAGTAATGCCTAGGAAATTGCCCTGATGTGGGGGATAACCATTGGAAACGATGGCTAATACCGCATAATACCTTCGGGTCAAAGAGGGGGACCTTCGGGCCTCTCGCGTCAGGATATGCCTAGGTGGGATTAGCTAGTTGGTGAGGTAATGGCTCACCAAGGCGACGATCCCTAGCTGGTCTGAGAGGATGATCAGCCACACTGGAACTGAGACACGGTCCAGACTCCTACGGGAGGCAGCAGTGGGGAATATTGCACAATGGGCGCAAGCCTGATGCAGCCATGCCGCGTGTGTGAAGAAGGCCTTCGGGTTGTAAAGCACTTTCAGTCGTGAGGAAGGTAGTGTAGTTAATAGCTGCATTATTTGACGTTAGCGACAGAAGAAGCACCGGCTAACTCCGTGCCAGCAGCCGCGGTAATACGGAGGGTGCGAGCGTTAATCGGAATTACTGGGCGTAAAGCGCATGCAGGTGGTTTGTTAAGTCAGATGTGAAAGCCCGGGGCTCAACCTCGGAATAGCATTTGAAACTGGCAGACTAGAGTACTGTAGAGGGGGGTAGAATTTCAGGTGTAGCGGTGAAATGCGTAGAGATCTGAAGGAATACCGGTGGCGAAGGCGGCCCCCTGGACAGATACTGACACTCAGATGCGAAAGCGTGGGGAGCAAACAGGATTAGATACCCTGGTAGTCCACGCCGTAAACGATGTCTACTTGGAGGTTGTGGCCTTGAGCCGTGGCTTTCGGAGCTAACGCGTTAAGTAGACCGCCTGGGGAGTACGGTCGCAAGATTAAAACTCAAATGAATTTGACGGGGGCCCGCACAAGCGGTGGAGCATGTGGTTTTAATTTCGATGCAACGCGAAGAACCTTACCTACTCTTGACATCCAGAGGACTTTGCAA

> *Bacillus cereus*

TGTACTGGCGGCGTGCCTATACTGCAAGTCGAGCGAATGGATTAAGAGCTTGCTCTTATGAAGTTAGCGGCGGACGGGTGAGTAACACGTGGGTAACCTGCCCATAAGACTGGGATAACTCCGGGAAACCGGGGCTAATACCGGATAACATTTTGAACCGCATGGTTCGAAATTGAAAGGCGGCTTCGGCTGTCACTTATGGATGGACCCGCGTCGCATTAGCTAGTTGGTGAGGTAACGGCTCACCAAGGCAACGATGCGTAGCCGACCTGAGAGGGTGATCGGCCACACTGGGACTGAGACACGGCCCAGACTCCTACGGGAGGCAGCAGTAGGGAATCTTCCGCAATGGACGAAAGTCTGACGGAGCAACGCCGCGTGAGTGATGAAGGCTTTCGGGTCGTAAAACTCTGTTGTTAGGGAAGAACAAGTGCTAGTTGAATAAGCTGGCACCTTGACGGTACCTAACCAGAAAGCCACGGCTAACTACGTGCCAGCAGCCGCGGTAATACGTAGGTGGCAAGCGTTATCCGGAATTATTGGGCGTAAAGCGCGCGCAGGTGGTTTCTTAAGTCTGATGTGAAAGCCCACGGCTCAACCGTGGAGGGTCATTGGAAACTGGGAGACTTGAGTGCAGAAGAGGAAAGTGGAATTCCATGTGTAGCGGTGAAATGCGTAGAGATATGGAGGAACACCAGTGGCGAAGGCGACTTTCTGGTCTGTAACTGACACTGAGGCGCGAAAGCGTGGGGAGCAAACAGGATTAGATACCCTGGTAGTCCACGCCGTAAACGATGAGTGCTAAGTGTTAGAGGGTTTCCGCCCTTTAGTGCTGAAGTTAACGCATTAAGCACTCCGCCTGGGGAGTACGGCCGCAAGGCTGAAACTCAAAGGAATTG

>*Pseudoalteromonas shioyasakiensis*

TACTGGGCGGCAGGTCTACCCTGCAAGTCGAGCGGAAACGAAGAGGTGCTTGCTCCTCTGGTGTCGAGCGGCGGACGGGTGAGTAATGCTTGGGAATATGCCTTATGGTGGGGGACAACAGTTGGAAACGACTGCTAATACCGCATGATGTCTACGGACCAAAGTGGGGGACCTTCGGGCCTCACGCCATAAGATTAGCCCAAGTGGGATTAGCTAGTTGGTGAGGTAATGGCTCACCAAGGCGACGATCCCTAGCTGGTTTGAGAGGATGATCAGCCACACTGGGACTGAGACACGGCCCAGACTCCTACGGGAGGCAGCAGTGGGGAATATTGCACAATGGGCGCAAGCCTGATGCAGCCATGCCGCGTGTGTGAAGAAGGCCTTCGGGTTGTAAAGCACTTTCAGTAAGGAGGAAAGGTTAAGTGTTAATAGCACTTAGCTGTGACGTTACTTACAGAAGAAGCACCGGCTAACTCCGTGCCAGCAGCCGCGGTAATACGGAGGGTGCGAGCGTTAATCGGAATTACTGGGCGTAAAGCGTACGCAGGCGGTTTGTTAAGCGAGATGTGAAAGCCCCGGGCTCAACCTGGGAACTGCATTTCGAACTGGCAAACTAGAGTGTGATAGAGGGTGGTAGAATTTCAGGTGTAGCGGTGAAATGCGTAGAGATCTGAAGGAATACCGATGGCGAAGGCAGCCACCTGGGTCAACACTGACGCTCATGTACGAAAGCGTGGGGAGCAAACAGGATTAGATACCCTGGTAGTCCACGCCGTAAACGATGTCTACTAGAAGCTCGGGTCTTCGGACTTGTTTTTCAAAGCTAACGCATTAAGTAGACCGCCTGGGGAGTACGGCCGCAAGGTTAAAACTCAAATGAATTGACGGGGGCCCGCACAAGCGGTGGAGCATGTGGTTTAATTCGATGCAACGCGAAGAACCTTACCTACACTGACATACAGAGAACTTACCAAAGATGGTTTGGTGCTTCGGGACCTCTGATACAGGTGCTGCATGGCTGTCGTCAGCTCGTGTTGTGGA

>*Vibrio rotiferianus*

CTACTGGCGGCAGCCTACCTGCAAGTCGAGCGGAAACGAGTTATCTGAACCTTCGGGGAACGATAACGGCGTCGAGCGGCGGACGGGTGAGTAATGCCTAGGAAATTGCCCTGATGTGGGGGATAACCATTGGAAACGATGGCTAATACCGCATAATGCCTACGGGCCAAAGAGGGGGACCTTCGGGCCTCTCGCGTCAGGATATGCCTAGGTGGGATTAGCTAGTTGGTGAGGTAATGGCTCACCAAGGCGACGATCCCTAGCTGGTCTGAGAGGATGATCAGCCACACTGGAACTGAGACACGGTCCAGACTCCTACGGGAGGCAGCAGTGGGGAATATTGCACAATGGGCGCAAGCCTGATGCAGCCATGCCGCGTGTGTGAAGAAGGCCTTCGGGTTGTAAAGCACTTTCAGTCGTGAGGAAGGTAGTGTAGTTAATAGCTGCATTATTTGACGTTAGCGACAGAAGAAGCACCGGCTAACTCCGTGCCAGCAGCCGCGGTAATACGGAGGGTGCGAGCGTTAATCGGAATTACTGGGCGTAAAGCGCATGCAGGTGGTTTGTTAAGTCAGATGTGAAAGCCCGGGGCTCAACCTCGGAATTGCATTTGAAACTGGCAGACTAGAGTACTGTAGAGGGGGGTAGAATTTCAGGTGTAGCGGTGAAATGCGTAGAGATCTGAAGGAATACCGGTGGCGAAGGCGGCCCCCTGGACAGATACTGACACTCAGATGCGAAAGCGTGGGGAGCAAACAGGATTAGATACCCTGGTAGTCCACGCCGTAAACGATGTCTACTTGGAGGTTGTGGCCTTGAGCCGTGGCTTTCGGAGCTAACGCGTTAAGTAGACCGCCCTGGGGAGTACGGTCGCAAGATTAAAACTCAAATGAATTGACGGGGGCCCGCACAAGCGTGAGCATGTGCTTTAATTCGATGCAACGCGAAGAATCTTACCTACTCTTGACATCCAGAGGAACTTTTCCAGAAGAATGGAA

> *Staphylococcus arlettae*

AGGTAACTGGCGGCGTGCCTATACTGCAAGTCGAGCGAACAGATAAGGAGCTTGCTCCTTTGACGTTAGCGGCGGACGGGTGAGTAACACGTGGGTAACCTACCTATAAGACTGGAATAACTCCGGGAAACCGGGGCTAATGCCGGATAACATTTAGAACCGCATGGTTCTAAAGTGAAAGATGGTTTTGCTATCACTTATAGATGGACCCGCGCCGTATTAGCTAGTTGGTAAGGTAATGGCTTACCAAGGCAACGATACGTAGCCGACCTGAGAGGGTGATCGGCCACACTGGAACTGAGACACGGTCCAGACTCCTACGGGAGGCAGCAGTAGGGAATCTTCCGCAATGGGCGAAAGCCTGACGGAGCAACGCCGCGTGAGTGATGAAGGGTTTCGGCTCGTAAAACTCTGTTATTAGGGAAGAACAAACGTGTAAGTAACTGTGCACGTCTTGACGGTACCTAATCAGAAAGCCACGGCTAACTACGTGCCAGCAGCCGCGGTAATACGTAGGTGGCAAGCGTTATCCGGAATTATTGGGCGTAAAGCGCGCGTAGGCGGTTTCTTAAGTCTGATGTGAAAGCCCACGGCTCAACCGTGGAGGGTCATTGGAAACTGGGAGACTTGAGTGCAGAAGAGGAAAGTGGAATTCCATGTGTAGCGGTGAAATGCGCAGAGATATGGAGGAACACCAGTGGCGAAGGCGACTTTCTGGTCTGTAACTGACGCTGATGTGCGAAAGCGTGGGGATCAAACAGGATTAGATACCCTGGTAGTCCACGCCGTAAACGATGAGTGCTAAGTGTTAAGGGGTTTCCGCCCCTTAGTGCTGCAGCTAACGCATTAAGCACTCCGCCTGGGGAGTACGACCGCAAGGTTGAAACTCAAAGGAATTGACGGGACCCGCACAAGCGGGAGCATGTGGTTTAATTCGAAGCACCCGAAGAACCTTACCAAATCTTGACATCCTTTGACCACTCT

>*Lysinibacillus fusiformis*

ACAACTGGCGGCGTGCCTATACTGCAGTCGAGCGAACAGAGAAGGAGCTTGCTCCTTTGACGTTAGCGGCGGACGGGTGAGTAACACGTGGGCAACCTACCCTATAGTTTGGGATAACTCCGGGAAACCGGGGCTAATACCGAATAATTTATTTCATCTCATGGTGAAATACTGAAAGACGGTTTCGGCTGTCGCTATAGGATGGGCCCGCGGCGCATTAGCTAGTTGGTGAGGTAACGGCTCACCAAGGCGACGATGCGTAGCCGACCTGAGAGGGTGATCGGCCACACTGGGACTGAGACACGGCCCAGACTCCTACGGGAGGCAGCAGTAGGGAATCTTCCACAATGGGCGAAAGCCTGATGGAGCAACGCCGCGTGAGTGAAGAAGGATTTCGGTTCGTAAAACTCTGTTGTAAGGGAAGAACAAGTACAGTAGTAACTGGCTGTACCTTGACGGTACCTTATTAGAAAGCCACGGCTAACTACGTGCCAGCAGCCGCGGTAATACGTAGGTGGCAAGCGTTGTCCGGAATTATTGGGCGTAAAGCGCGCGCAGGTGGTTTCTTAAGTCTGATGTGAAAGCCCACGGCTCAACCGTGGAGGGTCATTGGAAACTGGGAGACTTGAGTGCAGAAGAGGATAGTGGAATTCCAAGTGTAGCGGTGAAATGCGTAGAGATTTGGAGGAACACCAGTGGCGAAGGCGACTATCTGGTCTGTAACTGACACTGAGGCGCGAAAGCGTGGGGAGCAAACAGGATTAGATACCCTGGTAGTCCACGCCGTAAACGATGAGTGCTAAGTGTTAGGGGGTTTCCGCCCCTTAGTGCTGCAGCTAACGCATTAAGCACTCCGCCTGGGGAGTACGGTCGCAAGACTGAAACTCAAAGGAATTGACGGGGG
