## Supplementary Materials and Table 2 for "Parasitic Vectors in Aquaculture: *Neobenedenia girellae* and Leeches as Potential Transmission Agents for *Trypanosoma carassii spectrum* and Pathogenic bacteria": Supplementary file 3_Sequences of the Pathogens Isolated from Raoping.docx

>*Limnotrachelobdella okae*

TGGTCAACAAAACATAAAGATATTGGAACTTTATACTTCATTTTTGGTGCATGATCTGCAATATTAGGAACATCTATAAGATTTATTATTCGTTCAGAATTAGCTCAGCCAGGTAGTTTAATTGAAAATGATCAAATTTATAATTCTATAATTACAGCTCATGGTTTAATTATAATTTTCTTTATAGTTATACCTATTTTAATTGGTGGGTTTGGAAATTGATTAGTACCATTAATAATTGGGGCTCCAGACATAGCTTTTCCACGATTAAATAATCTAAGATTTTGGTTATTACCACCATCTTTAATTTTATTATTAGGATCAGCCTTAGTAGAAAGTGGAGTAGGGACAGGATGAACAGTTTATCCACCTTTATCTTCTAATATAGCTCACTCAGGTCCCTCAGTTGACATAGCAATCTTTTCTTTACATTTAGCAGGTGTATCTTCAATTCTAGGAGCTTTAAATTTTATTACAACAGTTGTTAATATACGATGAGAAGGTATAAAATTAGAACGTTTACCATTGTTTGTATGGGCAGTATTAATTACTGTGGTACTATTATTATTATCTTTACCAGTATTAGCTGCTGCTATTACTATATTATTAACAGATCGTAATTTAAATACAACATTTTTTGACCCTGTTGGTGGTGGAGATCCAATTTTATATCAACACTTATTTTGATTTTTTGGTCACCCTGGAAGTTTAA

>*Vibrio scophthalmi*

GGTACTGGGCGGCGGCTACCTGCAGTCGAGCGGTAACAGAGAGAAAGCTTGCTTTCTTTGCTGACGAGCGGCGGACGGGTGAGTAATGCCTGGGAATATGCCTTGATGTGGGGGATAACCATTGGAAACGATGGCTAATACCGCATAATGCCTACGGGCCAAAGAGGGGGATCTTCGGACCTCTCGCGTCAAGATTAGCCCAGGTGGGATTAGCTAGTTGGTGAGGTAATGGCTCACCAAGGCGACGATCCCTAGCTGGTCTGAGAGGATGATCAGCCACACTGGAACTGAGACACGGTCCAGACTCCTACGGGAGGCAGCAGTGGGGAATATTGCACAATGGGCGCAAGCCTGATGCAGCCATGCCGCGTGTGTGAAGAAGGCCTTCGGGTTGTAAAGCACTTTCAGTCGTGAGGAAGGTAGTTGCGTTAATAGCGTAATTATTTGACGTTAGCGACAGAAGAAGCACCGGCTAACTCCGTGCCAGCAGCCGCGGTAATACGGAGGGTGCGAGCGTTAATCGGAATTACTGGGCGTAAAGCGCATGCAGGTGGTTTGTTAAGTCAGATGTGAAAGCCCGGGGCTCAACCTCGGAATTGCATTTGAAACTGGCAGACTAGAGTACTGTAGAGGGGGGTAGAATTTCAGGTGTAGCGGTGAAATGCGTAGAGATCTGAAGGAATACCGGTGGCGAAGGCGGCCCCCTGGACAGATACTGACACTCAGATGCGAAAGCGTGGGGAGCAAACAGGATTAGATACCCTGGTAGTCCACGCCGTAAACGATGTCTACTTGGAGGTTGTGGCCTTGAGCCGTGGCTTTCGGAGCTAACGCGTTAAGTAGACCGCCTGGGGAGTACGGTCGCAAAATTAAAACTCAAATGAATTGACGGGGCCCCACAAGCGGTGGAGCATGTGGTTTAATTCGGATGCAACGCGAAGAACCTTACCTACTTCTTGACCTCCAGAAAAGCCAGCGGAAACCCAGG

>*Vibrio harveyi*

GTACCTGGGCGGCGGCCTACCTGCAAGTCGAGCGGAAACGAGTTATCTGAACCTTCGGGGAACGATAACGGCGTCGAGCGGCGGACGGGTGAGTAATGCCTAGGAAATTGCCCTGGTGTGGGGGATAACCATTGGAAACGATGGCTAATACCGCATAATACCTTCGGGTCAAAGAGGGGGACCTTCGGGCCTCTCGCGTCAGGATATGCCTAGGTGGGATTAGCTAGTTGGTGAGGTAATGGCTCACCAAGGCGACGATCCCTAGCTGGTCTGAGAGGATGATCAGCCACACTGGAACTGAGACACGGTCCAGACTCCTACGGGAGGCAGCAGTGGGGAATATTGCACAATGGGCGCAAGCCTGATGCAGCCATGCCGCGTGTGTGAAGAAGGCCTTCGGGTTGTAAAGCACTTTCAGTCGTGAGGAAGGTAGTGTAGTTAATAGCTGCATTATTTGACGTTAGCGACAGAAGAAGCACCGGCTAACTCCGTGCCAGCAGCCGCGGTAATACGGAGGGTGCGAGCGTTAATCGGAATTACTGGGCGTAAAGCGCATGCAGGTGGTTTGTTAAGTCAGATGTGAAAGCCCGGGGCTCAACCTCGGAATAGCATTTGAAACTGGCAGACTAGAGTACTGTAGAGGGGGGTAGAATTTCAGGTGTAGCGGTGAAATGCGTAGAGATCTGAAGGAATACCGGTGGCGAAGGCGGCCCCCTGGACAGATACTGACACTCAGATGCGAAAGCGTGGGGAGCAAACAGGATTAGATACCCTGGTAGTCCACGCCGTAAACGATGTCTACTTGGAGGTTGTGGCCTTGAGCCGTGGCTTTCGGAGCTAACGCGTTAAGTAGACCGCCTGGGGAGTACGGTCGCAAGATTAAAACTCAAATGAATTGACGGGGGCCCGCACAAGCGGTGGAGCATGTGGTTTAATTCGATGCAACCCGAAGAACCTTACCTACTCTGGACTTCCAAAAAACTTCCAAAGAATGGATTGGGTGCCTTCGGAACTCTGAGAACAGGTGCTGCAT

>*Photobacterium damselae*

GGACTGGCGGCGGCCTACCTGCAGTCGAGCGGAACTACTTACTGTTCCTTCTGGAACGTTAAGGGCGCCGATCGGCGATCGTTGAATAATGCCTGGGAATATGCCCTGGTGTGGGGGATAACTATTGGAAAACATACCTAATACCCCTTAATCTCTTCGAAGGGAACAGGGGGACCTTTCGGCCTCTCATTAGCCGATTTGGCAAGGTGGGGTTGGCTTGGTCTTGCATGACTGGCTCACCAAGGTGACGATCCCTAGCTGGGATCCGAGGATGATCAGCCACACTGACACTGATACTCCGCCCAGACTCCTACGGGAAGCTGAATCGGGGAATATTGCCCAATGGGGGAAACCCTGATGGAGCCATGACGGGTGTTTGAAGAAGGCCTTCTTTTTGCAAAGCACTTTACGAATGGAGGAAGGCCCGGAAACTAACACCTGCACTATTTGACGTTACCTACATAACAATCACCGGCTAACTCCGTGCCAGCAGACGCGGCAATACGGAGGGTGCAATCGTGAATCGGAATTACTGGGCGGAAAGCTCATGCAGGCGATCTGATATGCCAGATGTGAAACTCCGGACTGTAACCTCGGAAGTGAATTTGAAACTGGCAAGCTAGAGTCGTGTATAGAGGTGTAGAATTTCAGGTGAAACGCGAGAAATATCGAATAGATATGACGGAATACCAGTGGGTGAACGCGGACTCATGCTGACACACTGACGCTCAGATGCGAAAACCGTATGGAATTACAGGCTCTGCTACCCCTGCTACGTACACGTCGTATCGATGTCTACTTGGACGTTGTGGCCTTGATTCGTGACCTACGAGCTAACGCGATCGGCAGACGGCCTGGGGCGTACGATCCCAATATTAATGACTC

>*Micrococcus flavus*

AGGTGACTGGGCGGCGTGCTTACCTGCAAGTCGAACGATGAAGCCCAGCTTGCTGGGCGGATTAGTGGCGAACGGGTGAGTAACACGTGAGTAACCTGCCCTTGACTCTGGGATAAGCCCGGGAAACTGGGTCTAATACCGGATAGGAGCGCCTGTCGCATGACGGGTGCTGGAAAGATTTTTCGGTCATGGATGGACTCGCGGCCTATCAGCTTGTTGGTGAGGTAACGGCTCACCAAGGCGACGACGGGTAGCCGGCCTGAGAGGGTGACCGGCCACACTGGGACTGAGACACGGCCCAGACTCCTACGGGAGGCAGCAGTGGGGAATATTGCACAATGGGCGAAAGCCTGATGCAGCGACGCCGCGTGAGGGATGACGGCCTTCGGGTTGTAAACCTCTTTCAGCAGGGAAGAAGCGAAAGTGACGGTACCTGCAGAAGAAGCACCGGCTAACTACGTGCCAGCAGCCGCGGTAATACGTAGGGTGCGAGCGTTATCCGGAATTATTGGGCGTAAAGAGCTCGTAGGCGGTTTGTCACGTCTGTCGTGAAAGTCCGGGGCTTAACCCCGGATCTGCGGTGGGTACGGGCAGACTAGAGTGCAGTAGGGGAGACTGGAATTCCTGGTGTAGCGGTGGAATGCGCAGATATCAGGAGGAACACCGATGGCGAAGGCAGGTCTCTGGGCTGTAACTGACGCTGAGGAGCGAAAGCATGGGGAGCGAACAGGATTAGATACCCTGGTAGTCCATGCCGTAAACGTTGGGCACTAAGTGTGGGGACCATTCCACGGTTTCCGCGCCGCAGCTAACGCATTAAGTGCCCCGCCTGGGGAGTACGGCCGCAAGGCTAAAACTCAAAGGAATTGACGGGGGCCGCACAGCGGCGGAGCATGCGGATTAATTCGATGCAACGCGAAGAACCTTACCAAGGCTTGACATGTTTTCGACCGCCGGGGAAAACCCGGGG

>*Paracoccus zhejiangensis*

GGACTGGCGGCAGCCTAACCTGCAAGTCGAGCGAGACCTTCGGGTCTAGCGGCGGACGGGTGAGTAACGCGTGGGAATATGCCCTTCTCTACGGAATAGCCTCGGGAAACTGAGAGTAATACCGTATACGCCCTTCGGGGGAAAGATTTATCGGAGAAGGATTAGCCCGCGTTGGATTAGGTAGTTGGTGGGGTAATGGCCTACCAAGCCTACGATCCATAGCTGGTTTGAGAGGATGATCAGCCACACTGGGACTGAGACACGGCCCAGACTCCTACGGGAGGCAGCAGTGGGGAATCTTAGACAATGGGGGAAACCCTGATCTAGCCATGCCGCGTGAGTGATGAAGGCCTTAGGGTTGTAAAGCTCTTTCAGCTGGGAAGATAATGACGGTACCAGCAGAAGAAGCCCCGGCTAACTCCGTGCCAGCAGCCGCGGTAATACGGAGGGGGCTAGCGTTGTTCGGAATTACTGGGCGTAAAGCGCACGTAGGCGGACTGGAAAGTTGGAGGTGAAATCCCAGGGCTCAACCTTGGAACTGCCTTCAAAACTATCAGTCTGGAGTTCGAGAGAGGTGAGTGGAATACCGAGTGTAGAGGTGAAATTCGTAGATATTCGGTGGAACACCAGTGGCGAAGGCGGCTCACTGGCTCGATACTGACGCTGAGGTGCGAAAGCGTGGGGAGCAAACAGGATTAGATACCCTGGTAGTCCACGCCGTAAACGATGAATGCCAGACGTCGGGCAGCATGCTGTTCGGTGTCACACCTAACGGATTAAGCATTCCGCCTGGGGAGTACGGTCGCAAGATTAAAACTCAAAGGAATTGACGGGGGCCCGCACAAGCGGTGGAGCATGTGGTTTAATTCGAAGCAACGCGCAGAACCTTACCAACCCTTGACA

>*Priestia megaterium*

AGGTACCTGGCGGGTGCCTATACTGCAGTCGAGCGAACTGATTAGAAGCTTGCTTCTATGACGTTAGCGGCGGACGGGTGAGTAACACGTGGGCAACCTGCCTGTAAGACTGGGATAACTTCGGGAAACCGAAGCTAATACCGGATAGGATCTTCTCCTTCATGGGAGATGATTGAAAGATGGTTTCGGCTATCACTTACAGATGGGCCCGCGGTGCATTAGCTAGTTGGTGAGGTAACGGCTCACCAAGGCAACGATGCATAGCCGACCTGAGAGGGTGATCGGCCACACTGGGACTGAGACACGGCCCAGACTCCTACGGGAGGCAGCAGTAGGGAATCTTCCGCAATGGACGAAAGTCTGACGGAGCAACGCCGCGTGAGTGATGAAGGCTTTCGGGTCGTAAAACTCTGTTGTTAGGGAAGAACAAGTACGAGAGTAACTGCTCGTACCTTGACGGTACCTAACCAGAAAGCCACGGCTAACTACGTGCCAGCAGCCGCGGTAATACGTAGGTGGCAAGCGTTATCCGGAATTATTGGGCGTAAAGCGCGCGCAGGCGGTTTCTTAAGTCTGATGTGAAAGCCCACGGCTCAACCGTGGAGGGTCATTGGAAACTGGGGAACTTGAGTGCAGAAGAGAAAAGCGGAATTCCACGTGTAGCGGTGAAATGCGTAGAGATGTGGAGGAACACCAGTGGCGAAGGCGGCTTTTTGGTCTGTAACTGACGCTGAGGCGCGAAAGCGTGGGGAGCAAACAGGATTAGATACCCTGGTAGTCCACGCCGTAAACGATGAGTGCTAAGTGTTAGAGGGTTTCCGCCCTTTAGTGCTGCAGCTAACGCATTAAGCACTCCGCCTGGGGAGTACGGTC

>*Massilia suwonensis*

AGTGCGTGGGCGGCTGCTTTAACTGCAAGTCGAACGGCAGCGCGGGGCAACCTGGCGGCGAGTGGCGAACGGGTGAGTAATATATCGGAACGTACCCAAGAGTGGGGGATAACGTAGCGAAAGTTACGCTAATACCGCATACGATCTAAGGATGAAAGTGGGGGATCGCAAGACCTCATGCTCCTGGAGCGGCCGATATCTGATTAGCTAGTTGGTGAGGTAAAGGCTCACCAAGGCGACGATCAGTAGCTGGTCTGAGAGGACGACCAGCCACACTGGAACTGAGACACGGTCCAGACTCCTACGGGAGGCAGCAGTGGGGAATTTTGGACAATGGGCGCAAGCCTGATCCAGCAATGCCGCGTGAGTGAAGAAGGCCTTCGGGTTGTAAAGCTCTTTTGTCAGGGAAGAAACGGTAGAGGCTAATATCCTTTGCTAATGACGGTACCTGAAGAATAAGCACCGGCTAACTACGTGCCAGCAGCCGCGGTAATACGTAGGGTGCAAGCGTTAATCGGAATTACTGGGCGTAAAGCGTGCGCAGGCGGTTTTGTAAGTCTGTCGTGAAAGCCCCGGGCTCAACCTGGGAATTGCGATGGAGACTGCAAGGCTTGAATCTGGCAGAGGGGGGTAGAATTCCACGTGTAGCAGTGAAATGCGTAGAGATGTGGAGGAACACCGATGGCGAAGGCAGCCCCCTGGGTCAAGATTGACGCTCATGCACGAAAGCGTGGGGAGCAAACAGGATTAGATACCCTGGTAGTCCACGCCCTAAACGATGTCTACTAGTTGTCGGGTTTTAATTAACTTGGTCACGCAGCTAACGCGTGAAGTAGACCGCCTGGGGAGTACGGTCGCAAGATTAAAACTCAAGAATTGACGGGGGACCCGCACAAGCGGGGGATGATGTGGGATTAATTCGATGCAACGCGAAAAACCTTACCTACCCTTGACATGTCAGAAGTCCGGAGAAATCTGGATGTGGCCTCGAAAAGGGAACCTGGAACCACGGTGCCTGCATGCCTGCC

>*Vibrio marinisediminis*

TAGCTGGGCGGGCGGCCTAACCTGCAAGTCGAGCGGTAACAGGAAGAAAGCTTGCTTTCTTTGCTGACGAGCGGCGGACGGGTGAGTAATGCCTAGGAAATTGCCCTGATGTGGGGGATAACCATTGGAAACGATGGCTAATACCGCATAATCTCTTCGGAGCAAAGAGGGGGACCTTCGGGCCTCTCGCGTCAGGATATGCCTAGGTGGGATTAGCTAGTTGGTGAGGTAATGGCTCACCAAGGCGACGATCCCTAGCTGGTCTGAGAGGATGATCAGCCACACTGGAACTGAGACACGGTCCAGACTCCTACGGGAGGCAGCAGTGGGGAATATTGCACAATGGGCGCAAGCCTGATGCAGCCATGCCGCGTGTATGAAGAAGGCCTTCGGGTTGTAAAGTACTTTCAGTCGTGAGGAAGGCGTTAGTGTTAATAGCACTATCGTTTGACGTTAGCGACAGAAGAAGCACCGGCTAACTCCGTGCCAGCAGCCGCGGTAATACGGAGGGTGCGAGCGTTAATCGGAATTACTGGGCGTAAAGCGCATGCAGGTGGTTCGTTAAGTCAGATGTGAAAGCCCGGGGCTCAACCTCGGAATTGCATTTGAAACTGGCGGACTAGAGTACTGTAGAGGGGGGTAGAATTTCAGGTGTAGCGGTGAAATGCGTAGAGATCTGAAGGAATACCGGTGGCGAAGGCGGCCCCCTGGACAGATACTGACACTCAGATGCGAAAGCGTGGGGAGCAAACAGGATTAGATACCCTGGTAGTCCACGCCGTAAACGATGTCTACTTGGAGGTTGTGGCCTTGAGCCGTGGCTTTCGGAGCTAACGCGTTAAGTAGACCGCCTGGGGAGTACGGTCGCAAGATTAAAACTCAAATGAATTGACGGGGGCCCGCACAAGCGGGGGACATGGGGTTTAATTCGATGCACGCGAAGAACCTTACCTACTCTTGACCTCAGAAAAGCCCATAAAAGATACAGGTGGGGCCTTCCGGGCAA
