## Supplementary figures and images for "Parasitic Vectors in Aquaculture: *Neobenedenia girellae* and Leeches as Potential Transmission Agents for *Trypanosoma carassii spectrum* and Pathogenic bacteria"

### Fig. S1.tif

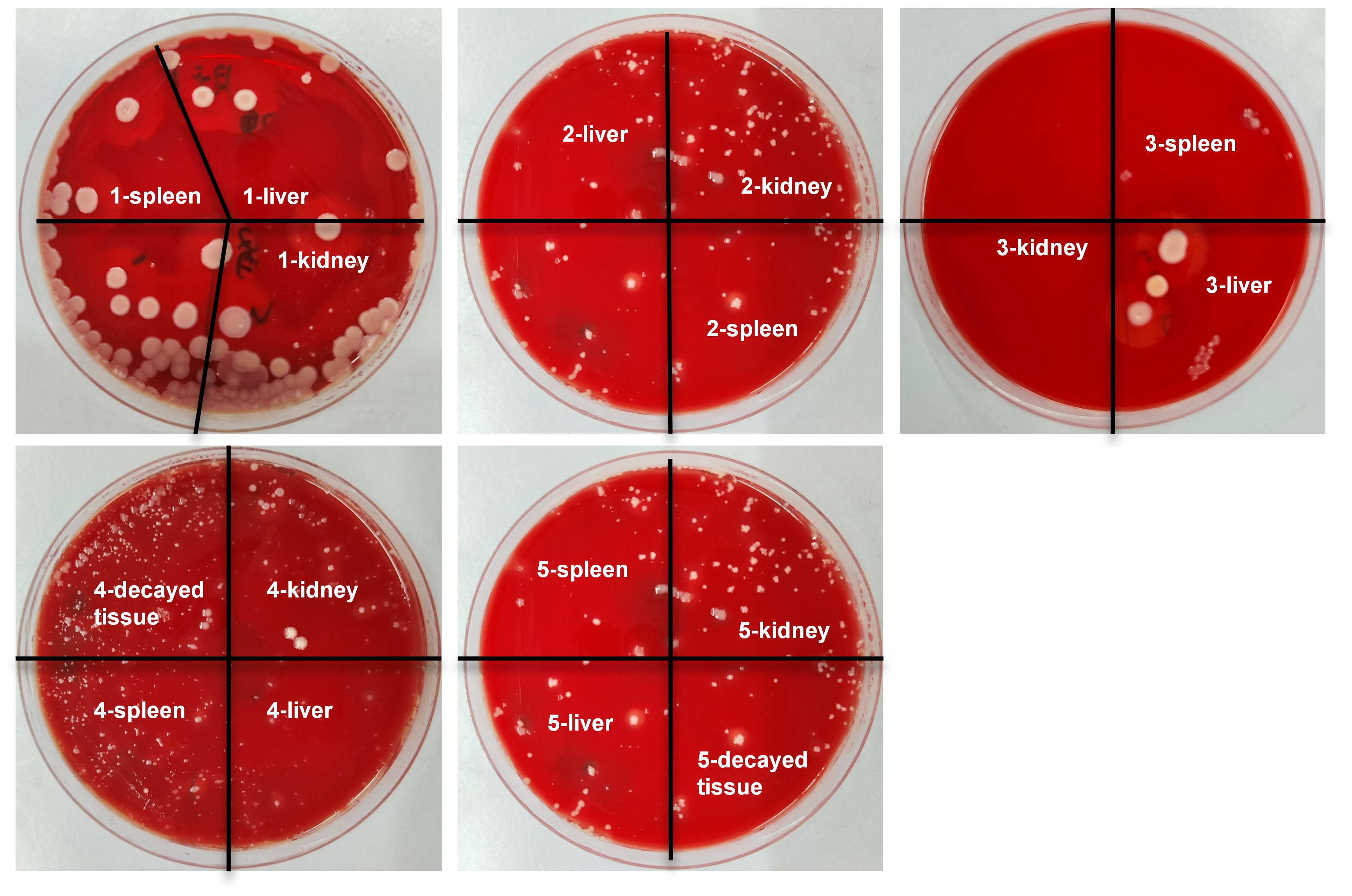

### Fig. S2.tif

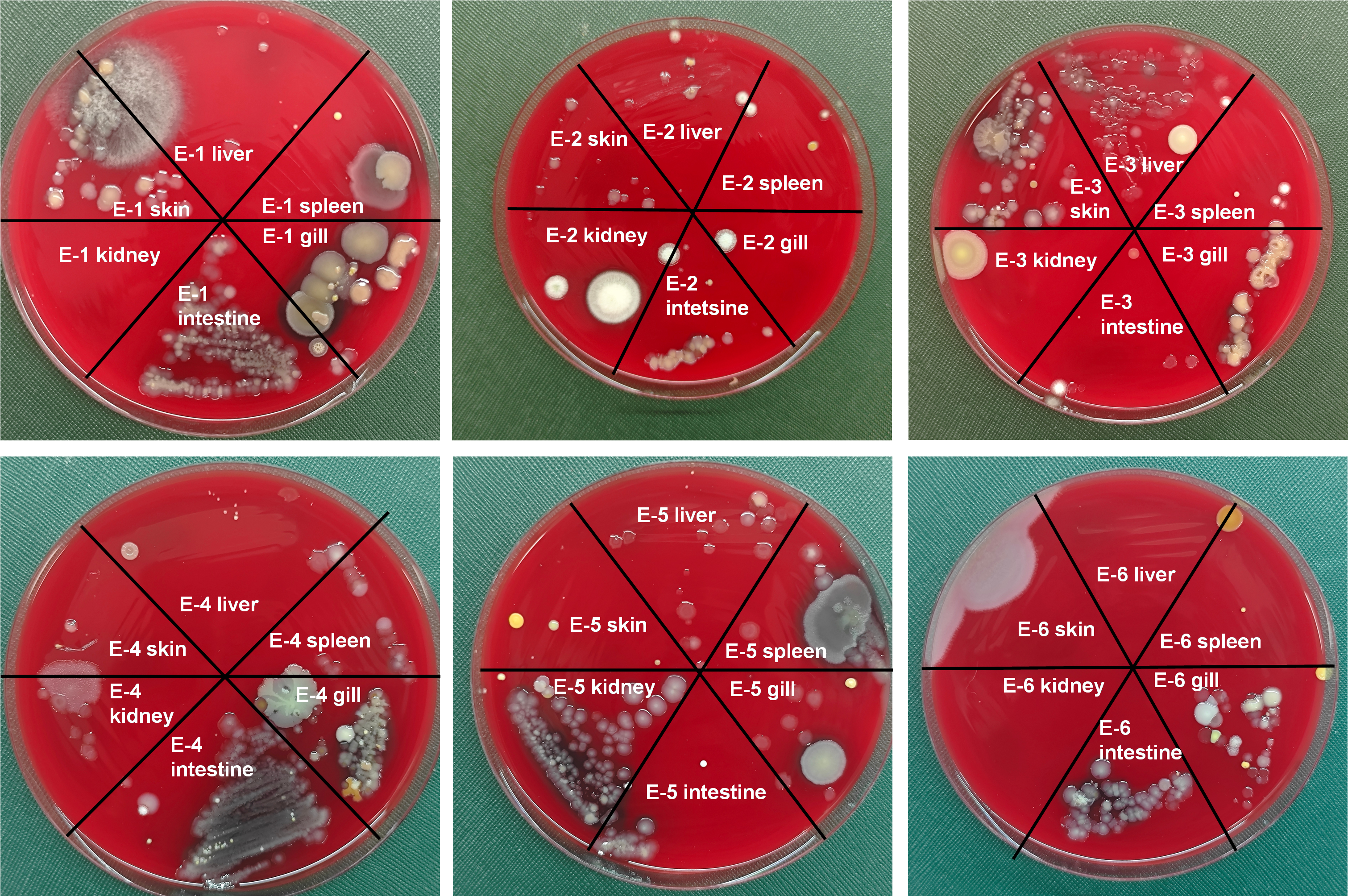

### Fig. S3.tif

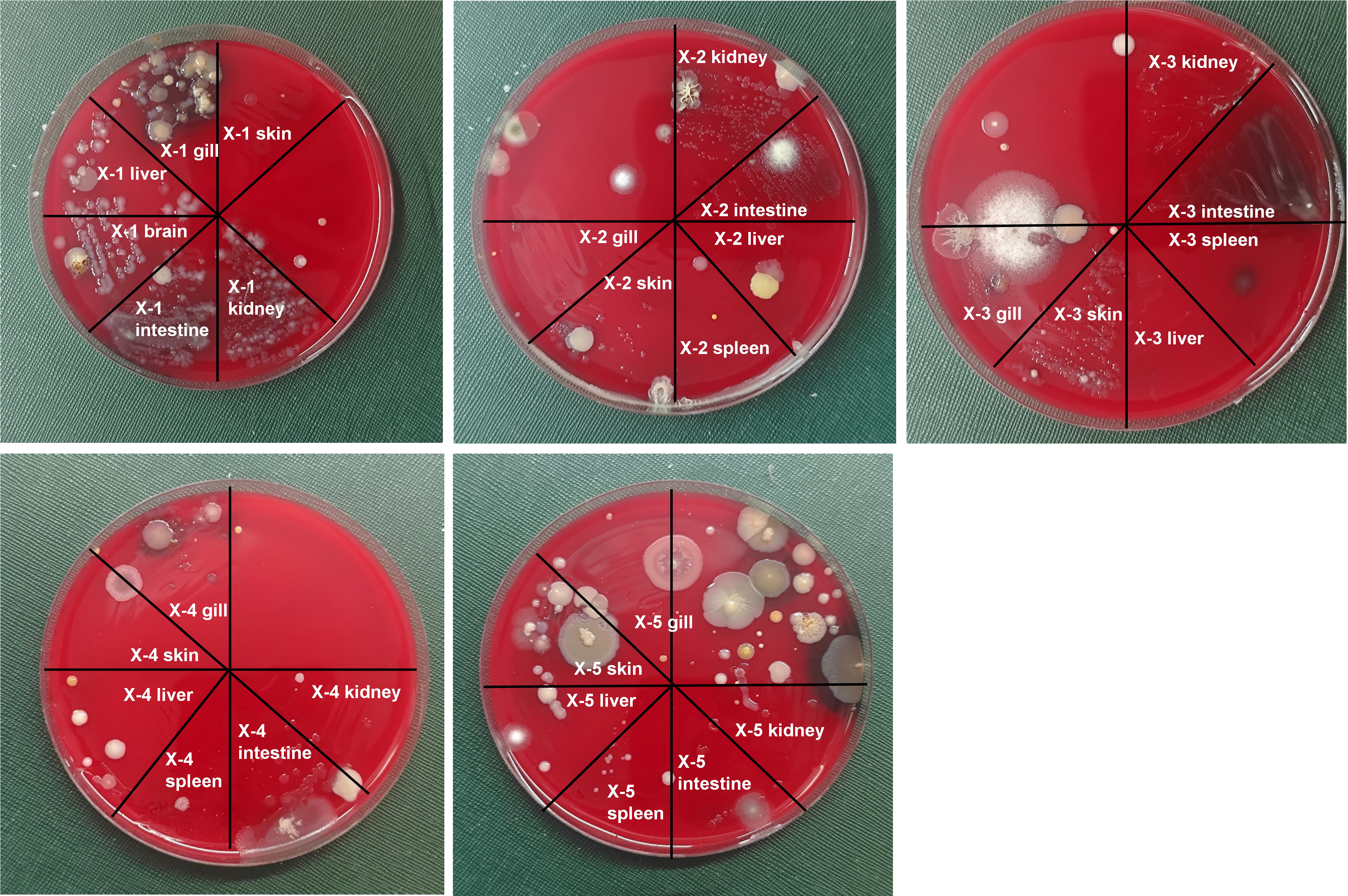
